# Myofibroblast transcriptome indicates SFRP2+ fibroblast progenitors in systemic sclerosis skin

**DOI:** 10.1101/2021.04.30.442148

**Authors:** Tracy Tabib, Mengqi Huang, Nina Morse, Anna Papazoglou, Rithika Behera, Minxue Jia, Melissa Bulik, Daisy E. Monier, Panayiotis V. Benos, Wei Chen, Robyn Domsic, Robert Lafyatis

**Affiliations:** Division of Rheumatology and Clinical Immunology, Department of Medicine, School of Medicine, University of Pittsburgh, Pittsburgh, PA, USA; Depatment of Computational and Systems Biology, School of Medicine, University of Pittsburgh, Pittsburgh, PA, USA; Division of Pulmonary Medicine, Allergy and Immunology, Department of Pediatrics, School of Medicine, University of Pittsburgh, Pittsburgh, PA, USA

## Abstract

Skin and lung fibrosis in systemic sclerosis (SSc) is driven by myofibroblasts, alpha-smooth muscle actin expressing cells that arise from a variety of cell types in murine fibrosis models. Utilizing single cell RNA-sequencing to examine the transcriptome changes, we show that SSc dermal myofibroblasts arise from an SFRP2/DPP4-expressing progenitor fibroblast population that globally upregulates expression of transcriptome markers, such as PRSS23 and THBS1. Only a fraction of SSc fibroblasts differentiate into myofibroblasts, as shown by expression of additional markers, SFRP4 and FNDC1. The myofibroblast transcriptome implicates upstream transcription factors that drive myofibroblast differentiation.

## INTRODUCTION

Skin fibrosis is a prominent clinical feature in most patients with systemic sclerosis (SSc, otherwise known as scleroderma), and the defining clinical feature for stratifying patients into two major disease subsets, limited or diffuse cutaneous disease. Skin tightness and thickening lead to considerable morbidity related mainly to contractures of hands as well as larger joints. It is also associated with pain, itching and cosmetic anguish^1^. Clinically skin involvement in SSc is associated with thickening, tethering, tightness and inflammation. Pathologically skin thickening is due to increased matrix deposition, most prominently type I collagen. Skin tightness may be due to this increase in matrix, but also correlates with the presence of myofibroblasts in the skin^2^. Thus, increased collagen production and the appearance of dermal myofibroblasts, typically seen first in the deep dermis, are pathogenic processes closely associated with the severity of clinical disease in SSc skin^3^.

In many fibrotic diseases myofibroblasts are the main collagen-producing cell driving fibrosis (reviewed in^4^). Perhaps more importantly in SSc skin, they exert tension on the tissue and through this mechanism may contribute to skin and joint contractures^2, 3, 5^. TGF-β and cell tension are factors most strongly implicated in myofibroblast development^6, 7^. Increasing matrix stiffness induces myofibroblast differentiation^8, 9^. TGF-β also induces myofibroblast differentiation and α-smooth muscle actin (SMA), the product of the ACTA2 gene and a robust though not specific marker of myofibroblasts in many different fibrotic diseases^10, 11^. Matrix stiffness and myofibroblast contraction also activate TGF-β^12, 13^, setting up a reinforcing amplification signal for tissue fibrosis. Several cytokines mediate, synergize with, or are permissive for the effect of TGF-β on myofibroblast formation: CTGF/CCN2^14, 15^, Endothelin-1^16^, and PDGF^17, 18, 19^. Others, such as FGF2, inhibit myofibroblast formation^20, 21^, while yet others, such as IFNγ, activate or inhibit myofibroblast formation in different fibrotic models^22, 23^.

Defining the phenotype of myofibroblasts beyond their expression of SMA has been challenging due to a rudimentary understanding of fibroblast heterogeneity in general and a paucity of specific markers of different fibroblast populations. However, recent studies have shed light on fibroblast heterogeneity in both mice and humans. In mice, markers are stable or dynamic (typically down regulated in adult mice) for dermal papilla (CRABP1), papillary (DPP4/CD26), and reticular (PDPN, SCA1/ATXN1) fibroblasts^24, 25^. Other investigators found that Engrailed/ DPP4-expressing fibroblasts in murine skin are profibrotic^26^. We have recently described two major and five minor fibroblast populations in normal skin^27^. The most common dermal fibroblast is long and slender, and expresses SFRP2 and DPP4. This population can be further divided into fibroblast subpopulations selectively expressing WIF1 and NKD2, or CD55 and PCOLCE2. A second major fibroblast population expresses FMO1 and MYOC. Minor fibroblast populations express CRABP1, COL11A1, FMO2, PRG4 or C2orf40. CRABP1-expressing fibroblasts most likely represent dermal papilla cells and COL11A1-expressing cells most likely represent dermal sheath cells, but FMO2, PRG4, C2orf40 minor fibroblast subsets are uncharacterized. Other recent studies have shown markers that distinguish between papillary (COL6A5, APCDD1, HSPB3, WIF1 and CD39) and reticular (CD36) dermal fibroblasts^28^.

Myofibroblasts are currently best defined by SMA staining, however SMA is expressed by a variety of other cell types including smooth muscle cells (SMCs), myoepithelial cells, pericytes and dermal sheath fibroblasts. These other cell types can be distinguished by expression of additional markers: Desmin (DES) and Smoothelin (SMTN) for SMCs^29^; Regulator of G Protein signaling 5 (RGS5), Chondroitin sulfate proteoglycan 4 CSPG4/NG2, Platelet-derived growth factor receptor beta (PDGFRB) for pericytes^30^; Keratin 5 (KRT5) and Keratin 14 (KRT14) for myoepithelial cells. However, there are no uniformly accepted specific markers for myofibroblasts. Cadherin 11 (CDH11) expression is associated with myofibroblast development and implicated in contractile force across myofibroblasts^31^, but CDH11 is expressed more diffusely by fibroblasts as well as by macrophages in SSc skin^32^. Thus, we lack specific markers for myofibroblasts and have limited understanding of the origins of this key pathogenic cell type in SSc skin.

The cellular progenitors of myofibroblasts in fibrotic disease models have been the source of increasingly sophisticated lineage tracing studies in mice, revealing that a variety of cell types can convert into myofibroblasts, including pericytes, epidermal cells, endothelial cells, preadipocytes, as well as fibroblasts (reviewed in^4^). In murine renal fibrosis, it appears that myofibroblasts arise from multiple progenitor cell types including resident fibroblasts and bone marrow derived cells, or transition from endothelial and epithelial cells^33^. Other lineage tracing studies have emphasized perivascular cells as progenitors of myofibroblasts in skin and muscle wound scarring^34^. In bleomycin-induced skin fibrosis adiponectin-expressing cells in adipose tissue can act as myofibroblast progenitors. In this model of SSc, transient cells co-expressing perilipin and SMA precede the development of myofibroblasts^35^. Although these studies suggest that multiple cell types can serve as progenitors of myofibroblasts in murine models, their origin in human disease, including SSc, has remained obscure. A recent study has shown that resident CD34+ fibroblasts in SSc skin undergo a change in phenotype characterized by downregulated CD34 expression and upregulated Podoplanin (PDPN) expression^36^. Although this phenotypic change was not strongly associated with the presence of myofibroblasts, markers of these cells were retained on myofibroblasts, suggesting that resident CD34+ dermal fibroblasts may be the precursors of myofibroblasts in SSc skin.

We here study the transcriptome-phenotypic changes of fibroblasts that occur in the skin from patients with SSc, focusing on identifying myofibroblasts using single cell RNA-sequencing (scRNA-seq) on total skin cell digests from SSc and healthy controls subjects.

## RESULTS

### Single cell transcriptomes from control skin

We have previously described fibroblast heterogeneity in normal skin^27^. We reanalyzed fibroblasts from normal healthy skin using an updated clustering algorithm and 4 additional discrete skin samples (Figures S1-S4). We observed the same cell populations we described previously based on the top differentially expressed genes (Table S1), but some additional populations were also apparent. SFRP2/DPP4-expressing fibroblasts, long narrow cells representing the most common population of fibroblasts, as before divided into two groups of cells: a WIF1/NKD2-expressing subgroup (cluster 1), also expressing HSPB3, APCDD1 and COL6A5, previously identified as markers of papillary dermis^28^, and a PCOLCE2/CD55/SLPI-expressing subgroup (cluster 0, Figure S1, S2, S3A-C). A second major population, expressing APOE, included MYOC/FMO1 fibroblasts described previously^27^, expressing low levels of APOE (APOE^lo^/MYOC, cluster 3), as well as a two subpopulations expressing higher levels of APOE, one of which also expressed high levels of C7 (APOE^hi^/C7, cluster 4), the other a APOE^hi^/C7 subset that also expressed high levels of CCL19, appearing mainly around vascular structures (APOE^hi^/C7/CCL19, cluster 7; Figures S3A-C). We identified several other cell populations based on previously described murine and human fibroblast markers surrounding hair follicles: CRABP1/COCH-expressing dermal papilla (cluster 5) and COL11A1/ACTA2-expressing dermal sheath cells (cluster 9, Figures S3A-C)25, 37. Two other cell populations cluster adjacent to dermal sheath and dermal papilla fibroblasts. One appears in the papillary dermis based on POSTN immunohistochemical staining (ASPN/POSTN, cluster 2, Figure S3C). The other expressed high levels of PTGDS (cluster 6), though this was not a specific marker. Finally, two small discrete populations of fibroblasts expressed SFRP4/ANGPTL7 (cluster 8) and SFRP4/LINC01133 (subset of cluster 3). We have previously described SFRP4-expressing fibroblasts in normal papillary dermis^38^. In our recent scRNA-seq description of normal fibroblast populations, we also described PRG4+ fibroblasts^27^. Although these cells did not form a discrete cluster on this reanalysis PRG4-expressing cells could be seen to group within PCOLCE2+ fibroblasts (not shown).

### Single cell transcriptomes from SSc and control skin

We then compared scRNA-seq of single cell suspensions from mid-forearm skin biopsies of 12 discrete samples from patients with SSc with the 10 control mid-forearm biopsy data described above. Control and SSc patients were balanced across sex (control=5/10 female; SSc=7/12 female), age (control mean age 51.9, median age = 57.5; SSc mean age =54.7, median age =57.5; Table 1). Similar numbers of cells were obtained from control (mean = 2821.6 cells/biopsy and median = 2623 cells/biopsy) and SSc (mean 3082 cells/biopsy and median = 3267 cells/biopsy). All patients with SSc had diffuse cutaneous disease with a mean MRSS = 26.1, median MRSS = 25. Disease duration was variable, between 0.48 and 6.48 years. Several of the patients were taking disease-modifying medications, as indicated (Table 1).

**Table 1:**
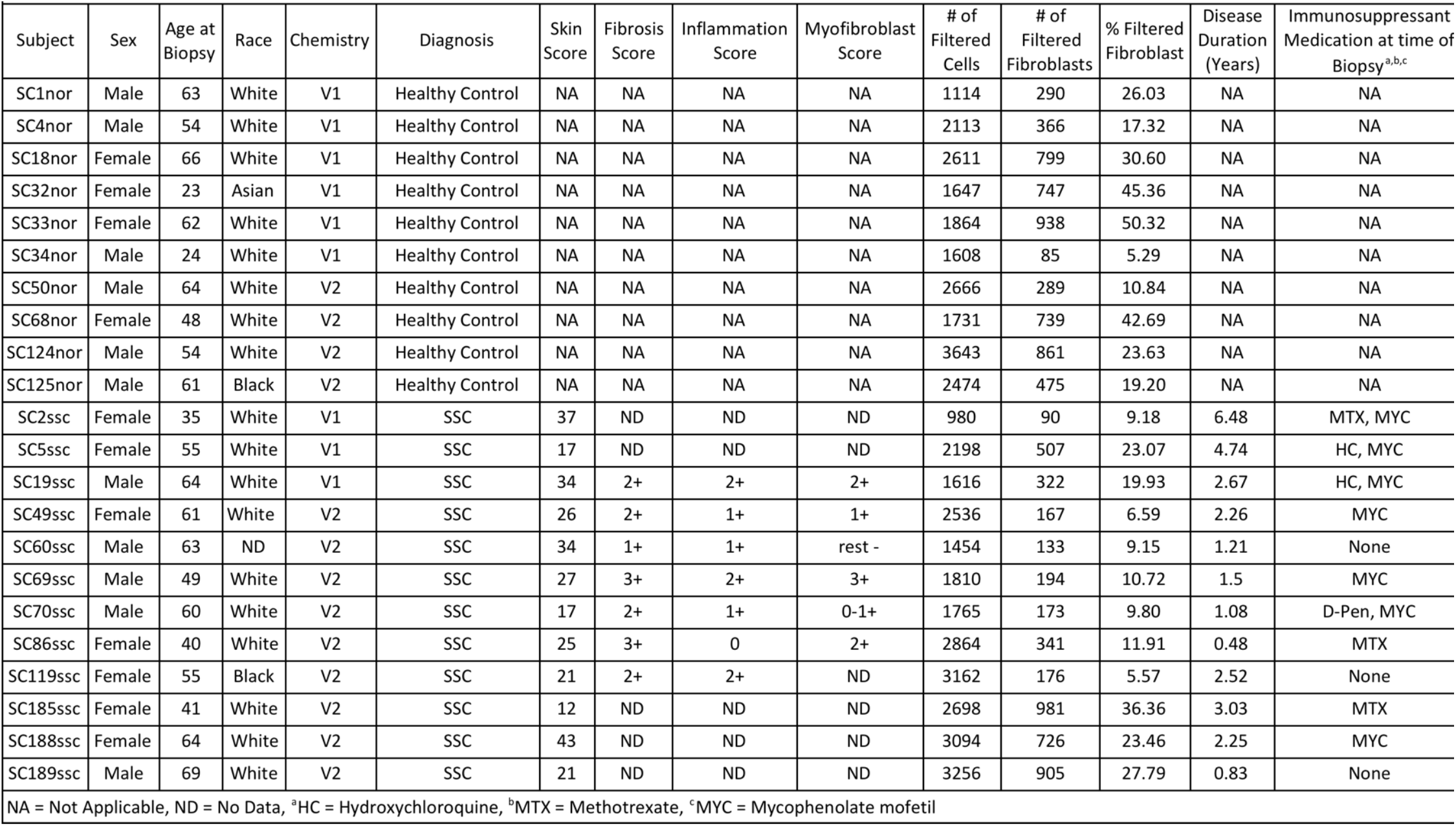
Subject Demographics, disease duration, medications and scRNA-seq fibroblast cell capture.

Cell transcriptomes were clustered by t-SNE, revealing all expected skin cell types, identified by examining the top differentially expressed genes in each cluster (Figure 1A, Table S2). Cell types in clusters were similar to cell types seen in normal skin (Figure 1A, S5A and S6^27^). Cells from each subject (Figure 1B) and chemistry (Figure S5B) were distributed in each cluster. The proportion of each cell population was generally preserved between healthy and SSc skin, however showing some changes in fibroblast and keratin 6A-expressing keratinocyte populations (Figure 1D). Notably, even at this low resolution, SSc fibroblasts can be seen to cluster separately from control fibroblasts, whereas for other cell types SSc and control cells largely overlie each other (Figure 1C). UMAP clustering of cells gave similar results (Figures S7-S9). We compared the average change in gene expression by SSc to healthy fibroblasts in each cluster (Table S3)

**Figure 1.**
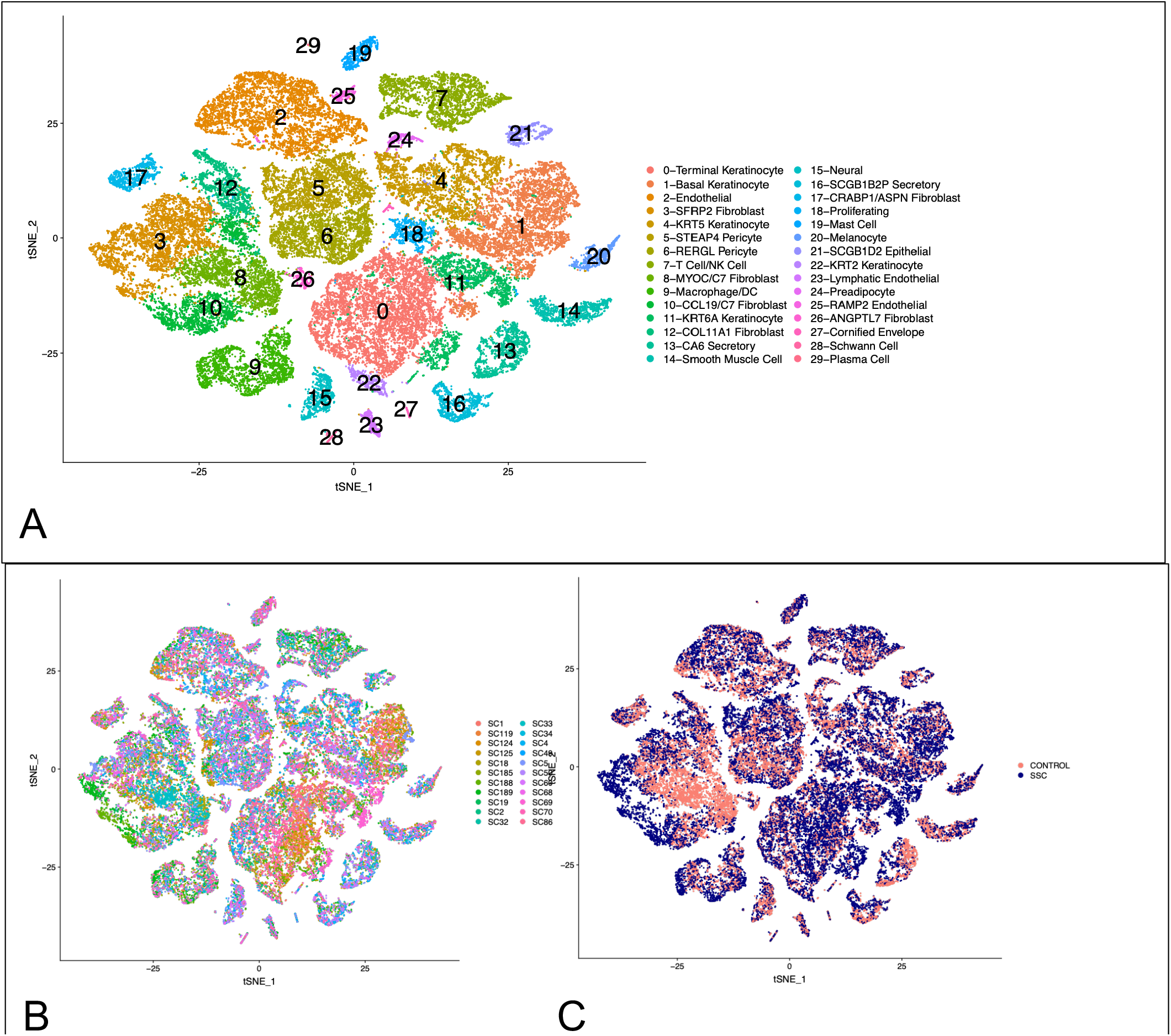
T-SNE plot of scRNA-seq data from control and SSc skin biopsies. Transcriptomes of all cells obtained after enzymatic digestion of dorsal mid-forearm skin biopsies from 10 healthy control and 12 SSc subjects, showing each SLM cluster by color (panel A) or by source from each patient (panel B) or by source form SSC (blue) or control (red) biopsies (panel C). Source data are provided as a Source Data file.

### Dermal fibroblast heterogeneity is preserved in SSc skin

We selected the cell clusters of fibroblasts based on expression of COL1A1, COL1A2 and PDGFRA (clusters 3, 8, 10, 12, 17 and 26 from Figure 1A), as we described previously these genes to be robust fibroblast cluster markers^27^. We reanalyzed just these cells by UMAP, revealing 10 fibroblast cell types (Figure 2A), generally paralleling those found in normal skin (see above and^27^). Fibroblast subclusters included cells from each subject (Figure 2B). However, fibroblasts from SSc patient skin samples showed prominent shifts between clusters (Figure 2C). Each subcluster could be identified by characteristic gene expression of top differentially expressed genes (Figures 2D and 3A, S10A and Table S4).

**Figure 2.**
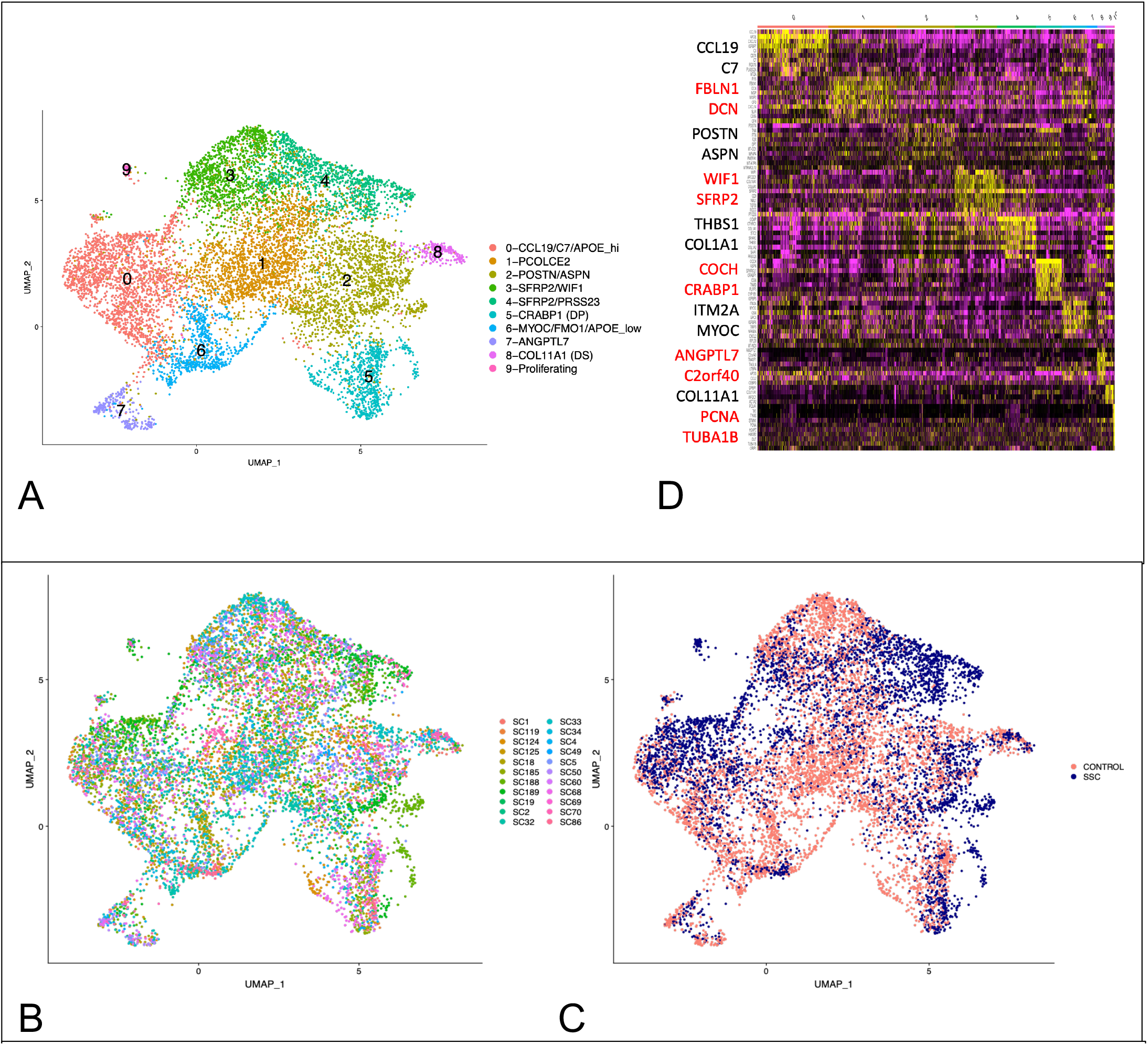
UMAP plot of scRNA-seq reclustering of fibroblasts and heatmap of subclusters. UMAP analysis of transcriptomes of fibroblasts (clusters 3, 8, 10, 12, 17 and 26 from Figure 1) from 10 healthy control and 12 SSc subjects, showing each SLM cluster by color (panel A) or by source from each patient (panel B) or by source form SSC (blue) or control (red) biopsies (panel C). Clustering of showing most differentially expressed genes associated with UMAP clusters. Yellow indicates increased expression, purple lower expression. Key marker genes are enlarged to the left (panel D).

**Figure 3.**
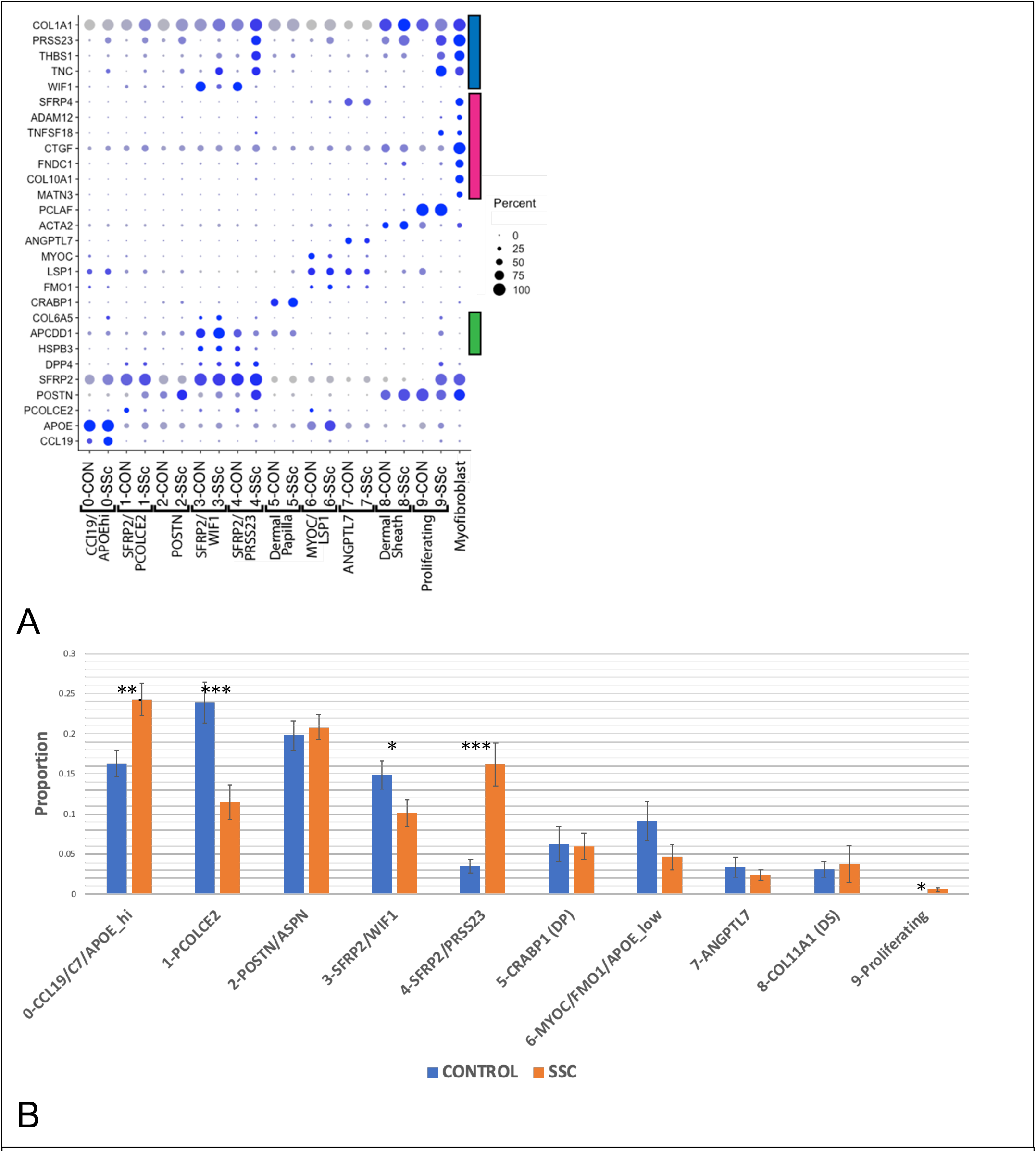
Key marker genes and proportions of fibroblast subclusters from control healthy (n=10) and SSc (n=12) skin. Dot plats of gene expression markers of fibroblasts populations in healthy and SSc skin (panel A). Subpopulations of fibroblasts including dermal sheath, dermal papilla, papillary (green bar), reticular and SSc fibroblasts (blue bar) and myofibroblasts (red bar) are indicated. Proportions of fibroblast subclusters as from control (blue) numbered and SSc biopsies (orange; clusters are numbered as in Figure 2 (panel B). Cell populations are differentially expressed between the groups (p<0.001, chi square test). Stars (*) indicate different proportions between SSc and Control subjects (p<0.05, error bars= SEM). Source data are provided as a Source Data file.

Fibroblasts expressing high levels of SFRP2 (SFRP2^hi^ fibroblasts, Figures 2A and 3A, fibroblast subclusters 1, 3 and 4), represent the major population of dermal fibroblasts, which are long slender cells found between collagen bundles^27^. SFRP2^hi^ fibroblasts included three subpopulations: subpopulations expressing WIF1 and NKD2 (WIF1+ fibroblasts, subcluster 3) and SLPI, PCOLCE2 and CD55 (PCOLCE2+ fibroblasts, subcluster 1) found previously in normal skin (Figures 2A and 3A), and a new subcluster of cells found mainly in SSc skin fibroblasts (PRSS23+ fibroblasts, subcluster 4; Figures 2A and 4A). Top and highly statistically significant GO terms associated with this new cluster were: Extracellular matrix organization and Extracellular structure organization (completely overlapping GO terms); Collagen fibril organization; Response to wounding; and Skeletal system development (Table S5).

**Figure 4.**
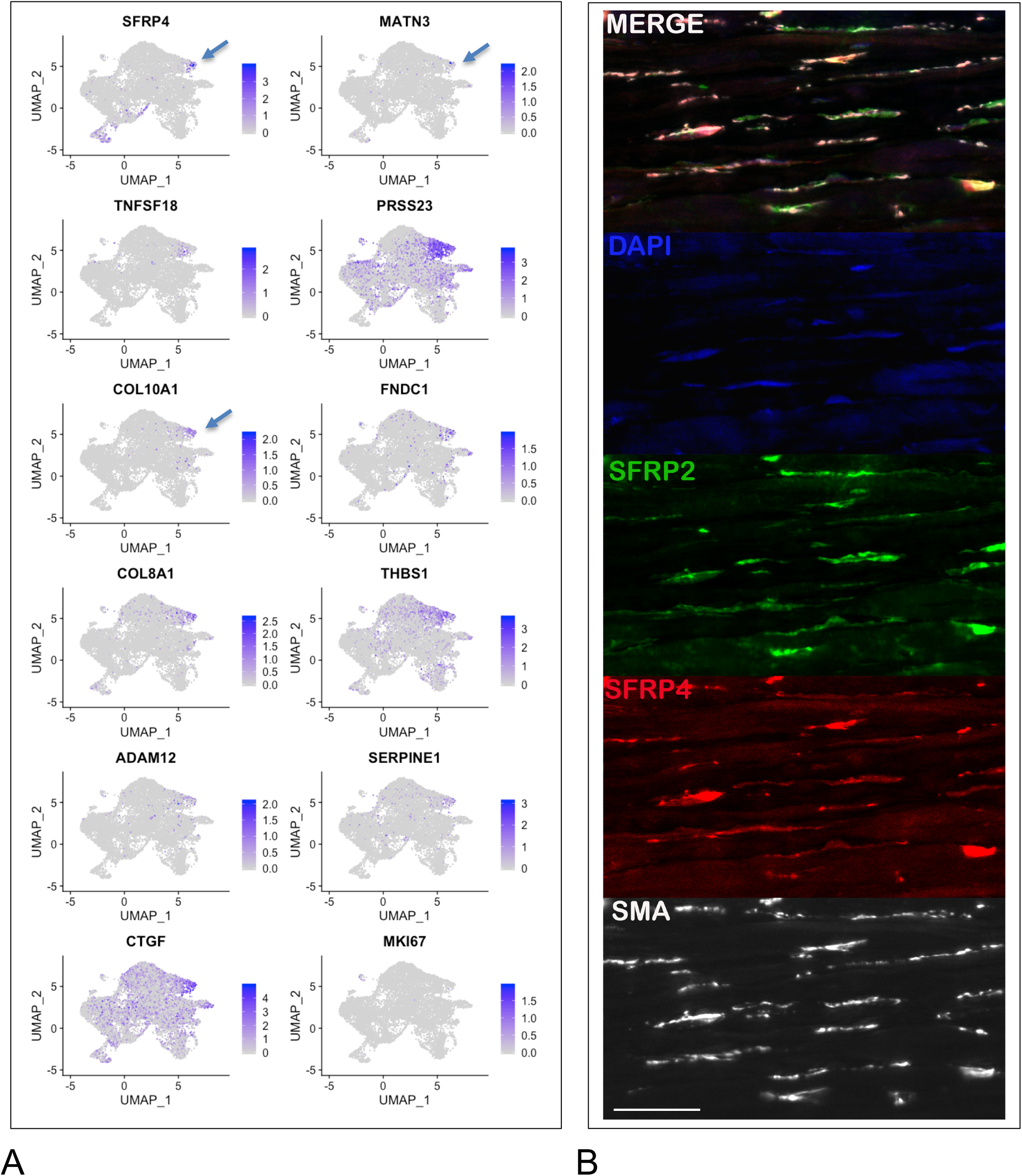
Feature plots and immunofluorescent staining of genes overexpressed by SFRP2+ SSc fibroblasts. PRSS23, THBS1, ADAM12, SERPINE1, FNDC1, CTGF show upregulated expression in cells associated more diffusely with fibroblast subcluster 4 (panel A). SFRP4, COL10A1, MATN3 show upregulated gene expression in a pattern more discrete within the cluster (arrows). Immunofluorescent staining of myofibroblasts in SSc skin (panel B). Deep dermis from a patient with diffuse cutaneous SSc co-stained with SFRP2 (green) and SFRP4 (red), SMA (white) show strong overlap between staining of myofibroblasts with SFRP2 and SFRP4. Nuclei (purple) are stained with Hoechst. Scale bar = 50 μM.

Expression of APOE defined cells in two clusters: APOE^hi^/CCL19/C7-expressing fibroblasts (clusters 0), and APOE^low^/FMO1/MYOC-expressing cells (subcluster 6, Figures 2A and 3A). We have previously identified this latter population fibroblast population as distributed in interstitial and perivascular regions^27^. The larger subpopulation of cells (subcluster 0) included a subgrouping of cells, highly expressing CCL19 showing a strong trend toward more SSc fibroblasts (Figure 3B), the SSc CCL19+ fibroblasts clustering separately from the control CCL19+ fibroblasts (Figure 2C), expressing higher levels of CCL19 (Figure 3A) and localizing primarily perivascularly (Figure S3C).

Three adjacent clusters showed markers of cells associated with hair follicles (subclusters 2, 5 and 8). CRABP1-expressing cells likely represent dermal papilla fibroblasts (DP, subcluster 5) and ACTA2/SOX2-expressing cells likely represent dermal sheath fibroblasts that may include dermal sheath stem cells (DS, cluster 8,^37, 39^). Cells in cluster 2 appear to represent cells closely related to hair follicles and/or papillary fibroblasts, as ASPN and F2R expressed by cells in this cluster stain brightly cells surrounding hair follicles (^40^ and see Human Atlas online), and POSTN stains brightly in the papillary matrix (Figure S3C). The close relationship between these cells is consistent with the observation that papillary dermal fibroblasts are required to regenerate hair follicles^24^. Other markers of papillary dermal cells^28^: APCDD1, HSPB3 and COL6A5 were expressed by cells in subcluster 3 (marked by a green bar in Figure 3A), part of the SFRP2-expressing population found also in the reticular dermis.

Our previous studies show SFRP4-staining fibroblasts in the papillary dermis of healthy as well as SSc skin^38^. Thus, ANGPTL7/C2orf40/SFRP4-expressing cells (subcluster 7, Figures 2A and 3A) represent a population of papillary fibroblasts, a second SFRP4+ population found only in SSc skin, characterized below, representing myofibroblasts (Figure 3A).

Collectively these studies indicate that the papillary dermis includes several different fibroblast populations, as they are found in subclusters 2, 3 and 7.

### SSc fibroblasts show global alterations in phenotype

Strikingly most of the fibroblasts from SSc patients clustered separately from the control subjects on UMAP dimensional reduction (Figure 2C). SSc fibroblasts clustered prominently in cluster 4 (SFRP2+/PRSS23+ fibroblasts) and also in a discrete region within cluster 0 (CCL19+ fibroblasts, Figure S10A). These two clusters showed proportionately more cells originating from SSc compared to healthy, control biopsies (Figure 3B). Reciprocal changes in cell proportions were seen in clusters 1, 3 and 6 with greater proportions of control cells in these clusters. In contrast to the marked separation between SSc and normal fibroblasts in the clusters above, fibroblasts predicted to reside in the papillary dermis, and DP and DS fibroblasts associated with hair follicles (subclusters 2, 5, and 8) were distributed in an approximately equal proportion between SSc and normal samples (Figures 2C and 3B). Together these results indicated a widespread shift in the phenotype of at least two different fibroblast populations in SSc reticular dermis, but not in fibroblast populations associated with the hair follicle and papillary dermis.

SFRP2^hi^/WIF1+ fibroblasts (subcluster 3) were largely depleted in SSc skin with the appearance of SFRP2 ^hi^ /PRSS23+ fibroblasts in the adjacent (subcluster 4). Comparing these two clusters directly revealed top differentially expressed genes including COMP and THBS1, genes highly associated with the MRSS and previously identified as biomarkers of skin disease^41, 42^ (Table S6).

### SSc fibroblasts show discrete altered gene expression

To investigate the changes in the transcriptome-phenotype of fibroblasts in SSc skin, we compared gene expression between SSc and control cells in each cluster (Table S7). Several of the highly upregulated genes in cluster 4 were recognizable as genes previously shown to correlate with the severity of SSc disease such as THBS1^41, 42^, TNC^43^, CTGF^42^, THY1^36^, CDH11^32^, and CCL2^44^ (Table 2). However, particularly striking to us was the marked upregulation of SFRP4, a gene we had studied previously in the context of a putative role for Wnts in SSc^38^. Further, on examining these and other genes increased in SSc SFRP2^hi^/PRSS23+ (Cluster 4) fibroblasts, we broadly observed two patterns of expression. Either genes were expressed by most cells in this cluster, such as PRSS23, THBS1 and TNC, or they were expressed by a subset of cells in this cluster, such as SFRP4, ADAM12, TNFSF18, CTGF, FNDC1, COL10A1, and MATN3 (Figures 3A and 4A). We did not see any suggestion of preadipocyte, pericyte or myeloid markers in myofibroblasts to suggest a transition from these cell types (Figure S10B).

**Table 2.**
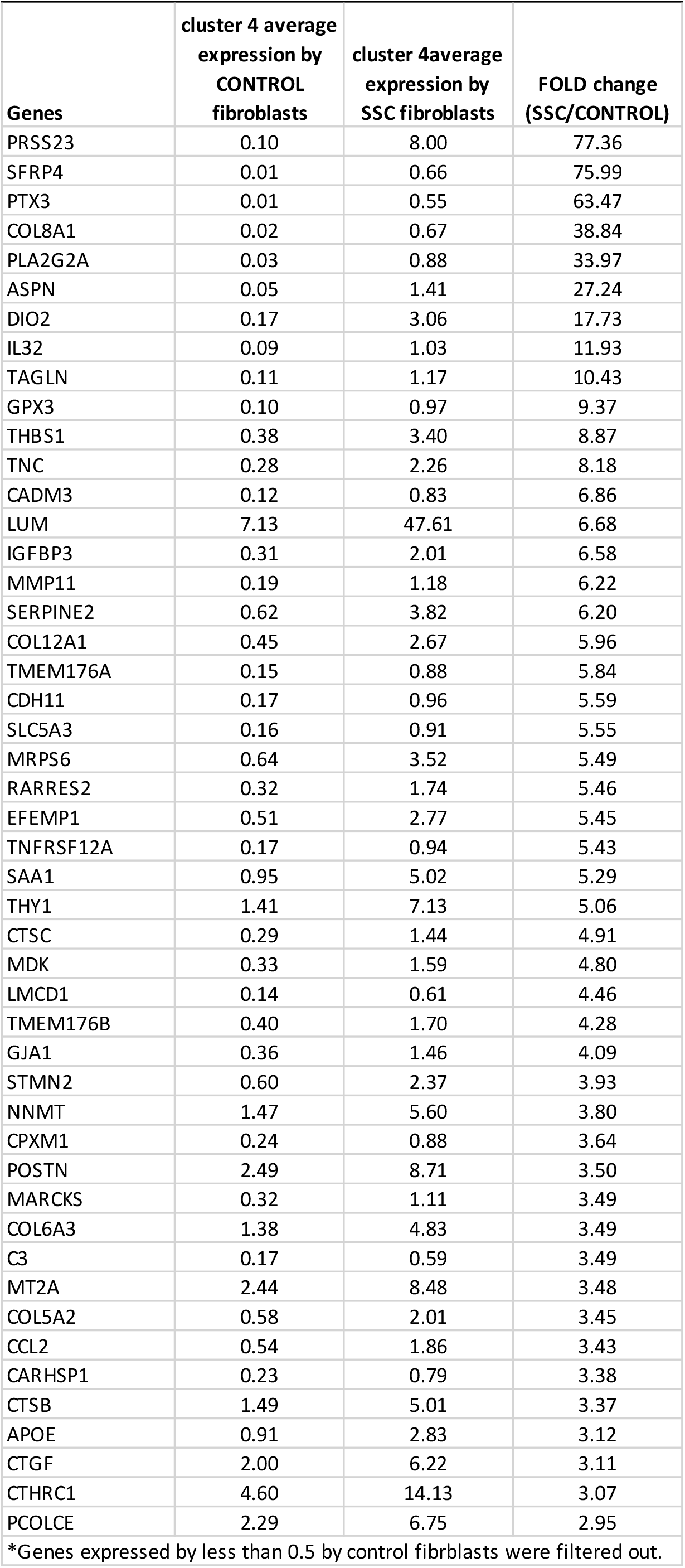
Upregulated gene expression by SSc fibroblasts in subcluster 4*.

### Myofibroblasts co-express SFRP2 and SFRP4

We showed previously that SFRP4 is upregulated in SSc skin and stains cells in the deep dermis, and that staining correlates with the MRSS^38^. However, at the time we did not associate this staining with myofibroblasts. Based on our scRNA-seq data showing a discrete cluster of SFRP2^hi^/SFRP4+ fibroblasts, we co-stained SFRP4 with SMA, the best-defined marker of myofibroblasts. We found that these two markers co-stain myofibroblasts in the deep dermis (Figures 4B, S10C). We have recently shown that SFRP2 stains long, thin cells in normal dermis^27^. Here, we show that SMA staining myofibroblasts co-stain with SFRP2, and that SFRP4-expressing cells also co-stain with SFRP2, indicating that SFRP2/SFRP4 co-expressing cells represent SSc dermal myofibroblasts.

### Myofibroblasts show a discrete transcriptome

We compared gene expression of SFRP2^hi^/PRSS23+/SFRP4-fibroblasts to SFRP2^hi^/PRSS23+/SFRP4+ myofibroblasts. The SFRP2^hi^/SFRP4+ fibroblasts were composed mostly of SSc cells (84/85 cells). Genes in addition to SFRP4 that are regulated (Table 3 and S8) included several other genes associated with the WNT pathway: SFRP1 and WNT2, and ACTA2, the gene encoding SMA (expressed 4.55-fold more highly in SFRP2^hi^SFRP4+ fibroblasts, Table S8).

**Table 3.**
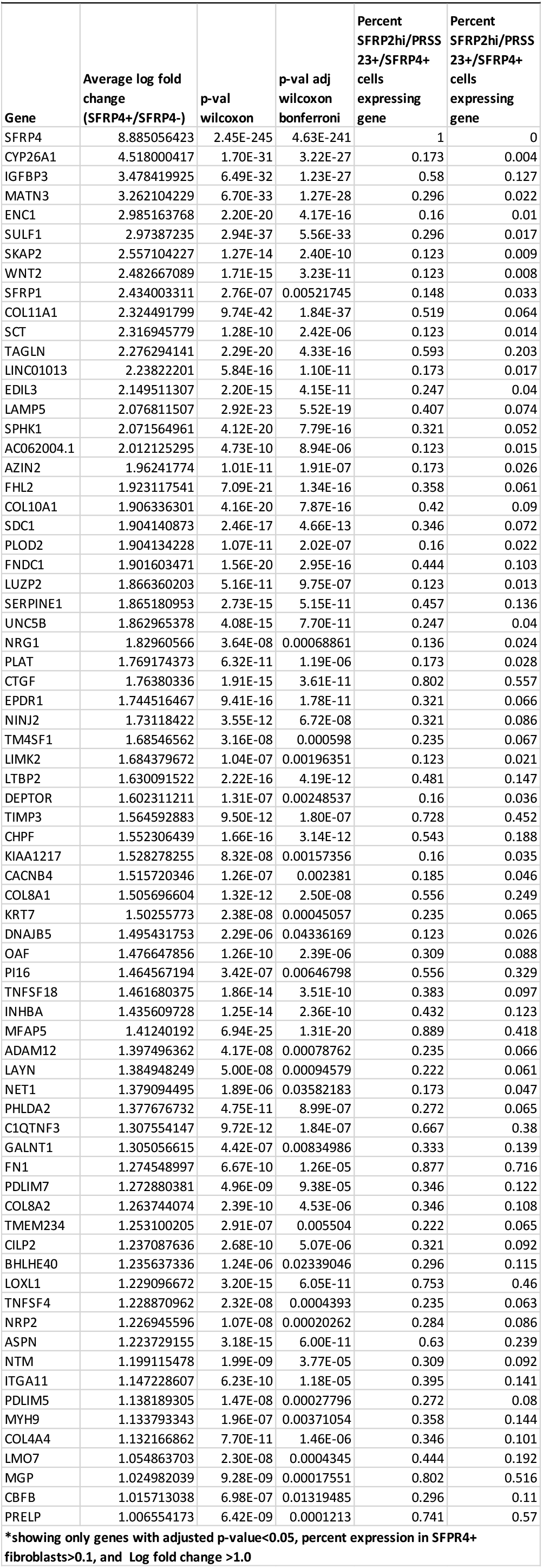
Gene expression by SSFRP2hi/PRSS23+/SFRP4+ myofibrobalsts compared to SFRP2^hi^/PRSS23+/SFRP4-fibroblasts*.

| Gene | Average log fold change (SFRP4+/SFRP4-) | p-val wilcoxon | p-val adj wilcoxon bonferroni | Percent SFRP2hi/PRSS 23+/SFRP4+ cells expressing gene | Percent SFRP2hi/PRSS 23+/SFRP4+ cells expressing gene |
| --- | --- | --- | --- | --- | --- |
| SFRP4 | 8.885056423 | 2.45E-245 | 4.63E-241 | 1 | 0 |
| CYP26A1 | 4.518000417 | 1.70E-31 | 3.22E-27 | 0.173 | 0.004 |
| IGFBP3 | 3.478419925 | 6.49E-32 | 1.23E-27 | 0.58 | 0.127 |
| MATN3 | 3.262104229 | 6.70E-33 | 1.27E-28 | 0.296 | 0.022 |
| ENC1 | 2.985163768 | 2.20E-20 | 4.17E-16 | 0.16 | 0.01 |
| SULF1 | 2.97387235 | 2.94E-37 | 5.56E-33 | 0.296 | 0.017 |
| SKAP2 | 2.557104227 | 1.27E-14 | 2.40E-10 | 0.123 | 0.009 |
| WNT2 | 2.482667089 | 1.71E-15 | 3.23E-11 | 0.123 | 0.008 |
| SFRP1 | 2.434003311 | 2.76E-07 | 0.00521745 | 0.148 | 0.033 |
| COL11A1 | 2.324491799 | 9.74E-42 | 1.84E-37 | 0.519 | 0.064 |
| SCT | 2.316945779 | 1.28E-10 | 2.42E-06 | 0.123 | 0.014 |
| TAGLN | 2.276294141 | 2.29E-20 | 4.33E-16 | 0.593 | 0.203 |
| LINC01013 | 2.23822201 | 5.84E-16 | 1.10E-11 | 0.173 | 0.017 |
| EDIL3 | 2.149511307 | 2.20E-15 | 4.15E-11 | 0.247 | 0.04 |
| LAMP5 | 2.076811507 | 2.92E-23 | 5.52E-19 | 0.407 | 0.074 |
| SPHK1 | 2.071564961 | 4.12E-20 | 7.79E-16 | 0.321 | 0.052 |
| AC062004.1 | 2.012125295 | 4.73E-10 | 8.94E-06 | 0.123 | 0.015 |
| AZIN2 | 1.96241774 | 1.01E-11 | 1.91E-07 | 0.173 | 0.026 |
| FHL2 | 1.923117541 | 7.09E-21 | 1.34E-16 | 0.358 | 0.061 |
| COL10A1 | 1.906336301 | 4.16E-20 | 7.87E-16 | 0.42 | 0.09 |
| SDC1 | 1.904140873 | 2.46E-17 | 4.66E-13 | 0.346 | 0.072 |
| PLOD2 | 1.904134228 | 1.07E-11 | 2.02E-07 | 0.16 | 0.022 |
| FNDC1 | 1.901603471 | 1.56E-20 | 2.95E-16 | 0.444 | 0.103 |
| LUZP2 | 1.866360203 | 5.16E-11 | 9.75E-07 | 0.123 | 0.013 |
| SERPINE1 | 1.865180953 | 2.73E-15 | 5.15E-11 | 0.457 | 0.136 |
| UNC5B | 1.862965378 | 4.08E-15 | 7.70E-11 | 0.247 | 0.04 |
| NRG1 | 1.82960566 | 3.64E-08 | 0.00068861 | 0.136 | 0.024 |
| PLAT | 1.769174373 | 6.32E-11 | 1.19E-06 | 0.173 | 0.028 |
| CTGF | 1.76380336 | 1.91E-15 | 3.61E-11 | 0.802 | 0.557 |
| EPDR1 | 1.744516467 | 9.41E-16 | 1.78E-11 | 0.321 | 0.066 |
| NINJ2 | 1.73118422 | 3.55E-12 | 6.72E-08 | 0.321 | 0.086 |
| TM4SF1 | 1.68546562 | 3.16E-08 | 0.000598 | 0.235 | 0.067 |
| LIMK2 | 1.684379672 | 1.04E-07 | 0.00196351 | 0.123 | 0.021 |
| LTBP2 | 1.630091522 | 2.22E-16 | 4.19E-12 | 0.481 | 0.147 |
| DEPTOR | 1.602311211 | 1.31E-07 | 0.00248537 | 0.16 | 0.036 |
| TIMP3 | 1.564592883 | 9.50E-12 | 1.80E-07 | 0.728 | 0.452 |
| CHPF | 1.552306439 | 1.66E-16 | 3.14E-12 | 0.543 | 0.188 |
| KIAA1217 | 1.528278255 | 8.32E-08 | 0.00157356 | 0.16 | 0.035 |
| CACNB4 | 1.515720346 | 1.26E-07 | 0.002381 | 0.185 | 0.046 |
| COL8A1 | 1.505696604 | 1.32E-12 | 2.50E-08 | 0.556 | 0.249 |
| KRT7 | 1.50255773 | 2.38E-08 | 0.00045057 | 0.235 | 0.065 |
| DNAJB5 | 1.495431753 | 2.29E-06 | 0.04336169 | 0.123 | 0.026 |
| OAF | 1.476647856 | 1.26E-10 | 2.39E-06 | 0.309 | 0.088 |
| PI16 | 1.464567194 | 3.42E-07 | 0.00646798 | 0.556 | 0.329 |
| TNFSF18 | 1.461680375 | 1.86E-14 | 3.51E-10 | 0.383 | 0.097 |
| INHBA | 1.435609728 | 1.25E-14 | 2.36E-10 | 0.432 | 0.123 |
| MFAP5 | 1.41240192 | 6.94E-25 | 1.31E-20 | 0.889 | 0.418 |
| ADAM12 | 1.397496362 | 4.17E-08 | 0.00078762 | 0.235 | 0.066 |
| LAYN | 1.384948249 | 5.00E-08 | 0.00094579 | 0.222 | 0.061 |
| NET1 | 1.379094495 | 1.89E-06 | 0.03582183 | 0.173 | 0.047 |
| PHLDA2 | 1.377676732 | 4.75E-11 | 8.99E-07 | 0.272 | 0.065 |
| C1QTNF3 | 1.307554147 | 9.72E-12 | 1.84E-07 | 0.667 | 0.38 |
| GALNT1 | 1.305056615 | 4.42E-07 | 0.00834986 | 0.333 | 0.139 |
| FN1 | 1.274548997 | 6.67E-10 | 1.26E-05 | 0.877 | 0.716 |
| PDLIM7 | 1.272880381 | 4.96E-09 | 9.38E-05 | 0.346 | 0.122 |
| COL8A2 | 1.263744074 | 2.39E-10 | 4.53E-06 | 0.346 | 0.108 |
| TMEM234 | 1.253100205 | 2.91E-07 | 0.005504 | 0.222 | 0.065 |
| CILP2 | 1.237087636 | 2.68E-10 | 5.07E-06 | 0.321 | 0.092 |
| BHLHE40 | 1.235637336 | 1.24E-06 | 0.02339046 | 0.296 | 0.115 |
| LOXL1 | 1.229096672 | 3.20E-15 | 6.05E-11 | 0.753 | 0.46 |
| TNFSF4 | 1.228870962 | 2.32E-08 | 0.0004393 | 0.235 | 0.063 |
| NRP2 | 1.226945596 | 1.07E-08 | 0.00020262 | 0.284 | 0.086 |
| ASPN | 1.223729155 | 3.18E-15 | 6.00E-11 | 0.63 | 0.239 |
| NTM | 1.199115478 | 1.99E-09 | 3.77E-05 | 0.309 | 0.092 |
| ITGA11 | 1.147228607 | 6.23E-10 | 1.18E-05 | 0.395 | 0.141 |
| PDLIM5 | 1.138189305 | 1.47E-08 | 0.00027796 | 0.272 | 0.08 |
| MYH9 | 1.133793343 | 1.96E-07 | 0.00371054 | 0.358 | 0.144 |
| COL4A4 | 1.132166862 | 7.70E-11 | 1.46E-06 | 0.346 | 0.101 |
| LMO7 | 1.054863703 | 2.30E-08 | 0.0004345 | 0.444 | 0.192 |
| MGP | 1.024982039 | 9.28E-09 | 0.00017551 | 0.802 | 0.516 |
| CBFB | 1.015713038 | 6.98E-07 | 0.01319485 | 0.296 | 0.11 |
| PRELP | 1.006554173 | 6.42E-09 | 0.0001213 | 0.741 | 0.57 |

\*showing only genes with adjusted p-value<0.05, percent expression in SFRP4+ fibroblasts>0.1, and Log fold change >1.0

### SFRP2^hi^WIF1+ fibroblasts are progenitors of myofibroblasts

To further investigate the relationship between SFRP2+ fibroblasts from healthy control skin, and fibroblasts and myofibroblasts in SSc skin, we used Monocle, an algorithm that tracks the relationship between single cell transcriptomes known as pseudotime ^45^. Pseudotime analysis indicated that there is a linear progression from SFRP2^hi^PCLOCE2+ fibroblasts (subcluster 1) to SFRP2^hi^WIF1+ fibroblasts (subcluster 3) to SFRP2^hi^PRSS23+WIF1− fibroblasts (subcluster 4) to SFRP2^hi^PRSS23+SFRP4+ myofibroblasts (Figure 5A and 5B). Although there is no polarity to the pseudotime analysis, since myofibroblasts are not present in normal skin, they most likely represent a later time in differentiation. This analysis indicates that SFRP2^hi^PRSS23+WIF1− fibroblasts are the immediate progenitors of myofibroblasts, and SFRP2^hi^WIF1+ fibroblasts the progenitors of SFRP2^hi^PRSS23+WIF1− fibroblasts.

**Figure 5.**
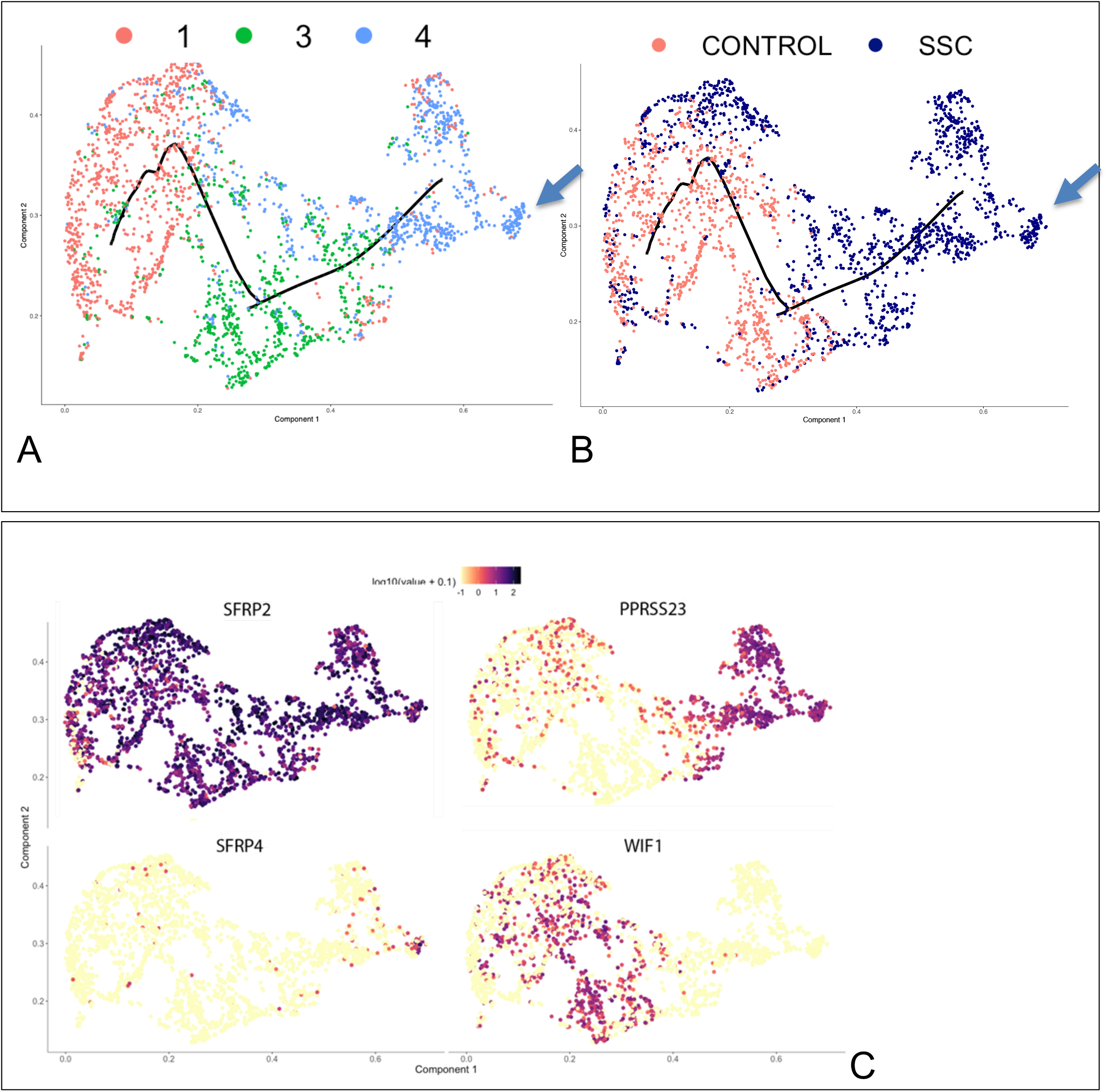
Pseudotime modeling of SFRP2+ fibroblast differentiation in SSc skin. Fibroblast subclusters defined in Figure 2 were analyzed using Monocle with the trajectory as indicated by the black line, with cells colored by subcluster of origin: subclsuter 1 (red), subcluster 3 (green) and subcluster 4 (blue; panel A) or by subject status: healthy control (red) and SSc (blue; panel B). SFRP2 was expressed by all of the cells (panel C, right lower panel). PRSS23 was expressed more highly by cells clustered later in pseudotime, corresponding to fibroblast subcluster 4 (panel C). SFRP4 was expressed more highly expressed even later in pseudotime, corresponding to myofibroblasts identified in t-SNE plots (Figures 3A and 4A).

This analysis reinforced the upregulated gene expression, transcriptome markers identified by examining the transcriptomes of SFRP2^hi^SFRP4+ myofibroblasts. These included COL10A1, FNDC1, SERPINE1, MATN3 and CTGF (Figure S11).

To further support the trajectory of SFRP2-expressing fibroblasts, we applied Velocyto, analyzing the single-cell RNA seq data based on spliced and unspliced transcript reads ^46^, supporting movement of SFRP2^hi^WIF1+ to SFRP2^hi^PRSS23+WIF1− fibroblasts to SFRP2^hi^PRSS23+SFRP4+ fibroblasts (Figure S12).

### Increased proliferating SFRP2^hi^PRSS23+WIF1− fibroblasts in SSc skin

A minor population of fibroblasts (subcluster 9), clustered separately from the other fibroblasts because they highly differentially expressed genes associated with cell proliferation (including PCNA and PCLAF, Figure 6A). We have shown previously that macrophages in IPF lungs expressing these markers are indeed proliferating cells^47^, though in this case these are extremely rare (representing only 0.38 % of the fibroblasts and .080 % of the total cells) and thus unlikely to be detected by immunohistochemistry. Of the 39 cells, only 2 of the cells in this subcluster were from healthy skin. The other 37 cells were from the SSc skin samples. The 2 cells from healthy skin showed markers of dermal sheath cells (DPEP1 and COL11A1, Figure 6B). In contrast, all of the cells from the SSc patients expressed markers of SFRP2^hi^PRSS23+WIF1− cells (SFRP2, PRSS23, TNC, COL10A1). However, these cells did not selectively express markers associated with differentiation of SFRP2^hi^PRSS23+WIF1− cells into myofibroblasts (not shown).

**Figure 6.**
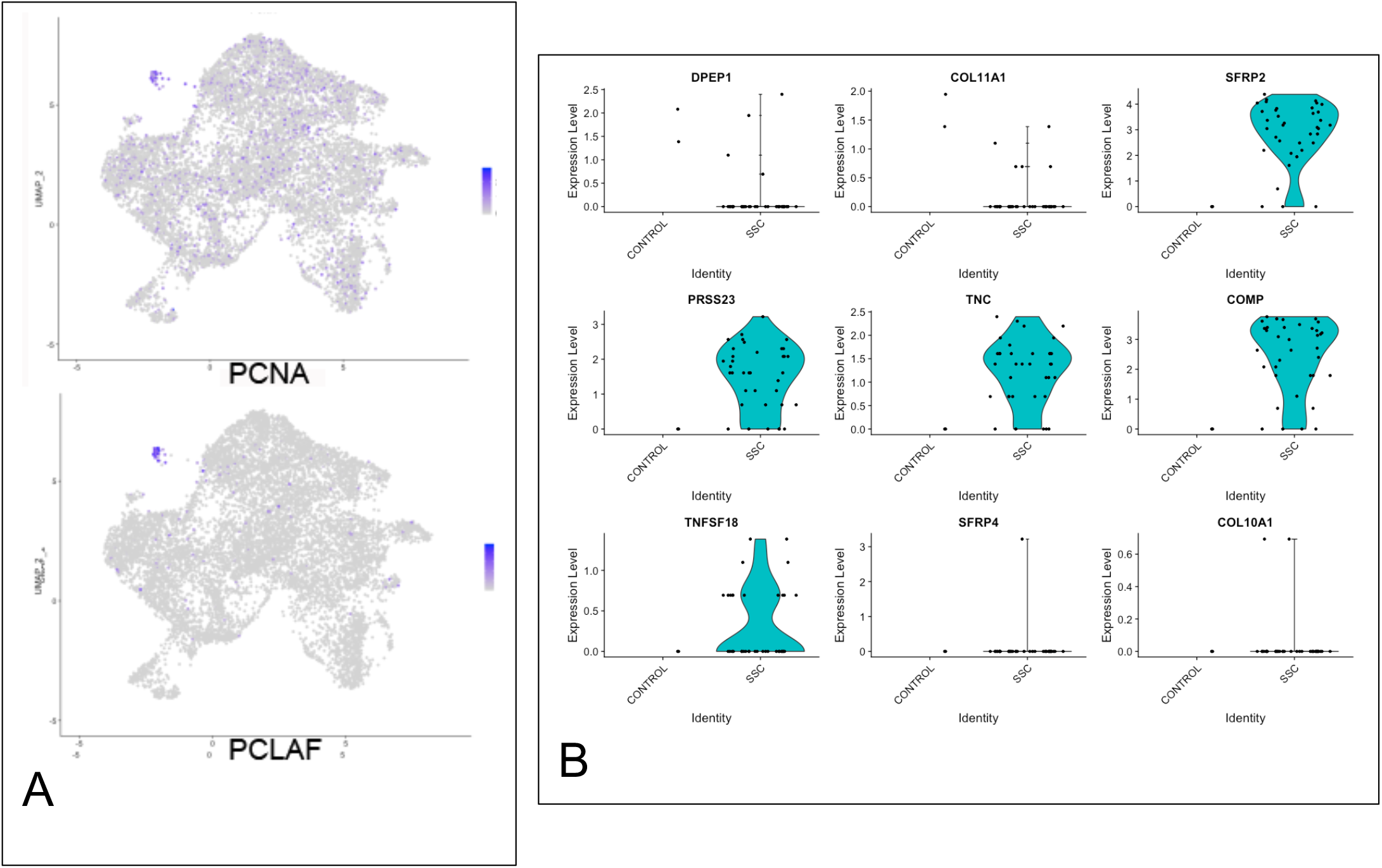
Proliferating fibroblasts in healthy control and SSc skin. Feature plots indicating that expression of proliferation markers, PCNA and PCLAF, are limited to cells in subcluster 9 (panel A). Violin plots indicate gene expression by proliferating cells (subcluster 9), showing markers of dermal sheath cells (DPEP1 and COL11A1) by healthy control cells (2 cells) and markers of SFRP2hiPRSS23+WIF1− cells (SFRP2, PRSS23, TNC, COMP and TNFSF18) by SSc fibroblasts (37 cells). Only one SSc fibroblast expressed markers of myofibroblasts (SFRP4 and COL10A1).

### Correlation between bulk microarray and SFRP2+/SFRP4+ cells

Several previous studies have examined bulk mRNA expression in SSc skin^42, 48, 49, 50^. Analyzing microarray data from our previous biomarker study and clinical trials conducted by our center using the same microarray platform^42, 51, 52, 53^, we compared genes upregulated in SSc whole biopsy gene expression data with our single cell results. Several microarray clusters showing genes upregulated in subsets of SSc patients contained genes expressed more highly by SSc fibroblasts or myofibroblasts, emphasizing the important role these genes play in these signatures (Figure 7A). Probing the single cell dataset with these clusters as gene modules showed that, indeed, they detect the global change in SSc fibroblasts (PRSS23 signature), or the change associated with myofibroblasts (SFRP4 signature) or (COL10A1 signature, Figure 7B). We have shown in previous publications that expression of several of the genes in these clusters (THBS1, COMP, ADAM12, and CTGF) correlate highly and statistically significantly with the MRSS^42^, so the observation that these genes cluster together in bulk RNA-seq analysis and their co-expression in our single cell RNA-seq dataset in the transition of healthy SFPR2+ fibroblasts to SSc SFRP2+ fibroblasts (THBS1) and myofibroblasts (ADAM12 and CTGF) is consistent with the roles of these SSc fibroblast populations in driving clinical disease. Expression of PRSS23, a marker for the first step in SSc fibroblast differentiation, correlated highly with the MRSS (Figure 7C), confirming that the first step in SSc fibroblast differentiation is associated with clinical skin disease.

**Figure 7.**
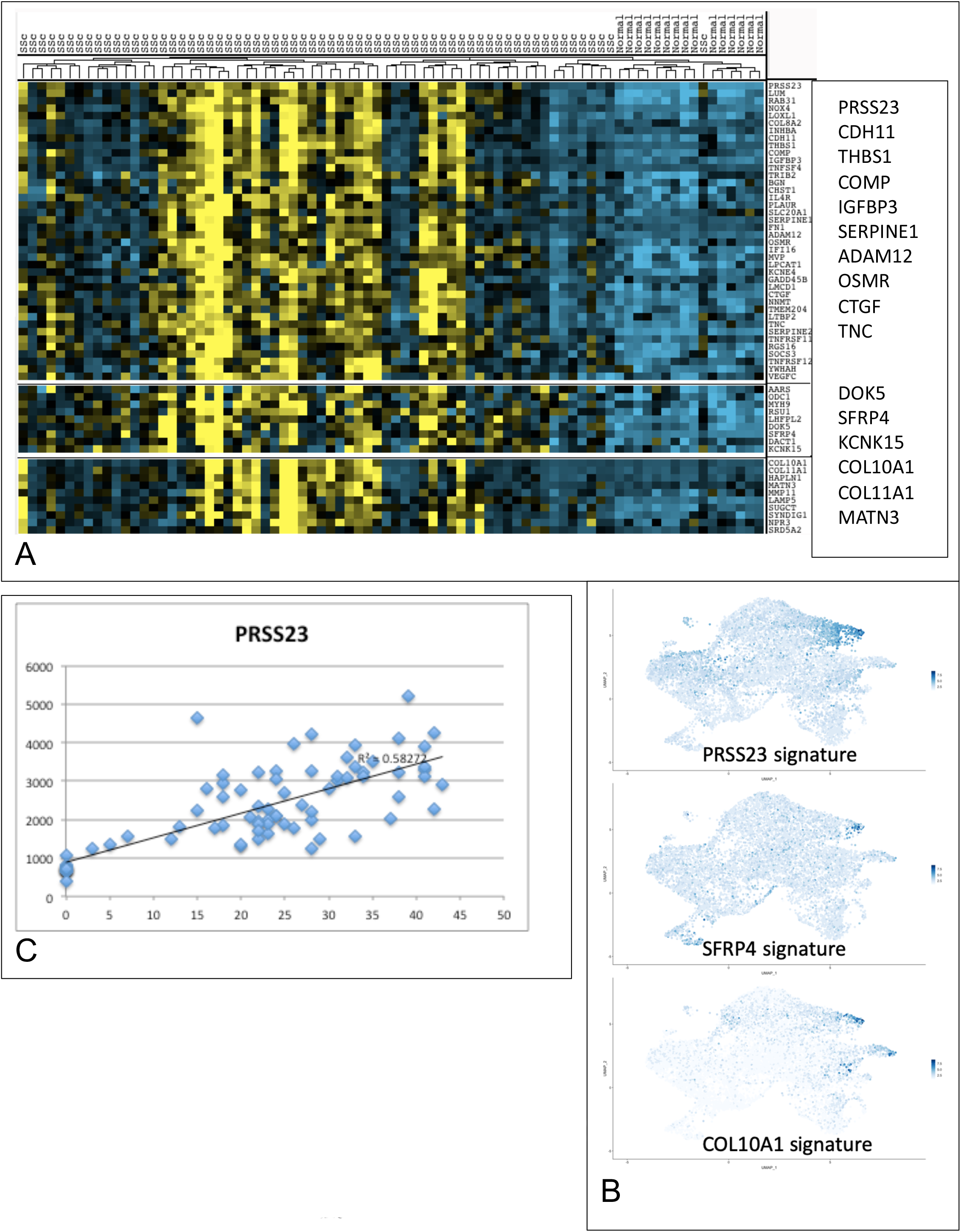
Bulk RNA expression data clusters reflect gene expression by SSc fibroblasts and myofibroblasts. Bulk gene expression from microarrays of patients with SSc (n=66) and healthy control skin (n=9) as indicated clustered hierarchically (panel A; yellow=high, blue= low expression). Feature plots corresponding to gene signature in of each cluster are shown (panel B, Seurat AddModuleScore function). PRSS23 expression on microarray correlates highly with the MRSS (R2=0.58272, panel C).

### Predicted transcription factor regulation of SSc fibroblast and myofibroblast gene expression

A significant challenge to gaining further insight into disease pathogenesis is relating gene expression changes to underlying alterations in intracellular signaling. To address this question, we analyzed our data using SCENIC, a computational method developed for detecting transcription factors (TF) networks^54^. T-SNE analysis of subclusters 1-4 by regulon (rather than by gene) showed a clear separation of the SSc fibroblasts from subcluster 4 (Figures 8A and 8B). Clustering the TFs driving regulons plots revealed a series of TFs, including TGIF2, FOSL2, RUNX1, STAT1 and IRF7 (Figure 8C and 8D), these genes expressed more highly also in this cell population (Figure 8E).

**Figure 8.**
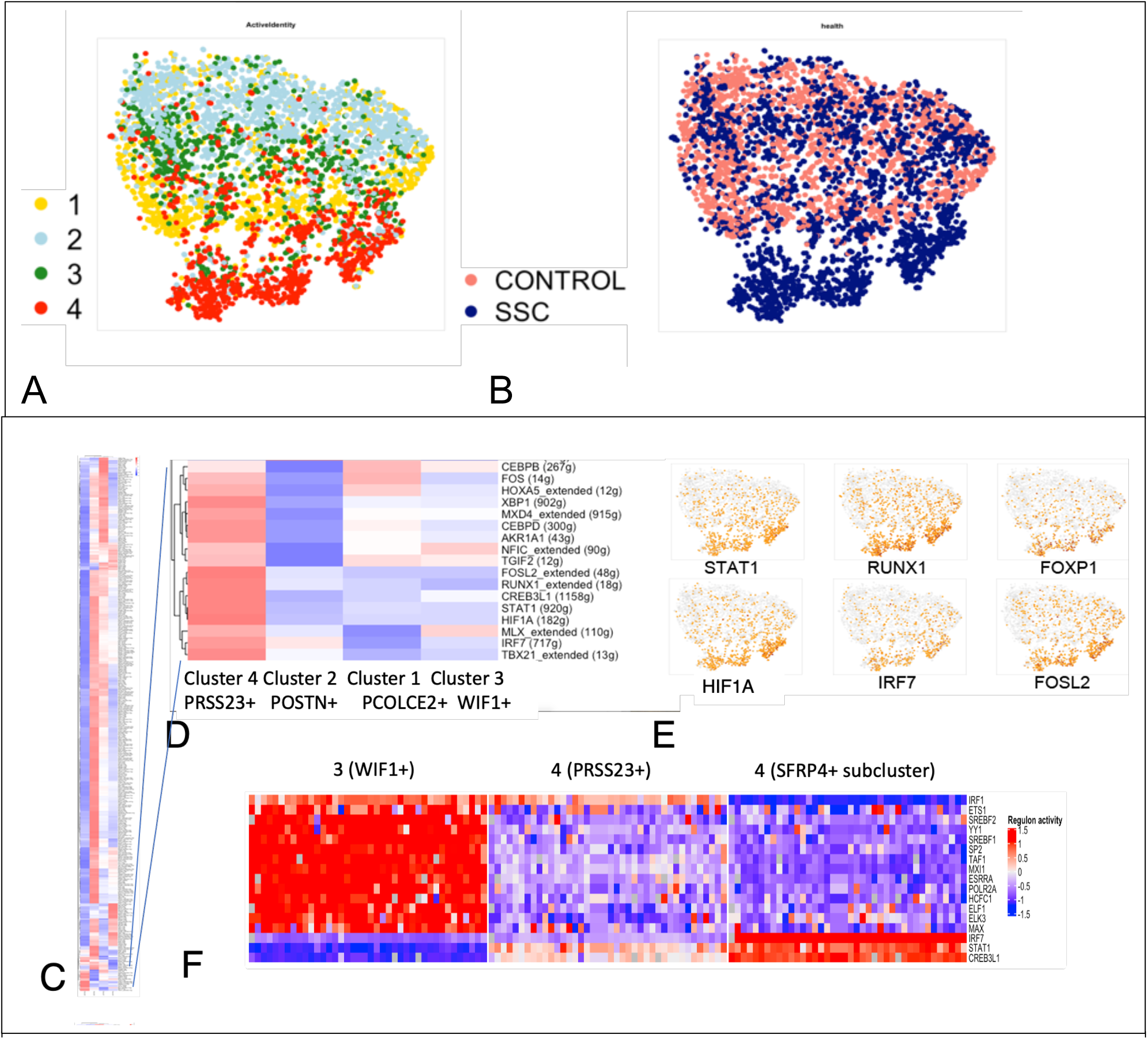
Regulons associated with TFs. Clustering of fibroblast subclusters 1-4 by regulon expression, colored according to gene expression subclusters as in Figure 2 and indicated in the legend (panel A) and according to SSc/healthy disease status (panel B). Clustering of regulons identified comparing fibroblast subclusters 1-4 (panel C), expanded section showing regulons upregulated in cluster 4 (panel D). Gene expression indicated on T-SNE plots of select TFs (panel E, brown=increased expression). Heatmap showing iterative downsamplings and SCENIC analysis of regulon activities comparing SFRP2+WIF1+ cluster 3 (WIF1+) with subcluster 4 divided into myofibroblasts (SFRP4+) and SFRP2+PRSS23+WIF1− non-myofibroblasts (PRSS23+, Panel F). Regulons associated with 38 non-redundant out of 40 SCENIC analyses are shown. For clustering panels, red=high, blue=low expression.

To examine TFs associated with myofibroblast differentiation more selectively, we divided the SFRP2+WIF1−SFRP4+ (myofibroblast populations) from the SFRP2+PRSS23+ (remainder of subcluster 4, composed mainly of SSc fibroblasts) and compared the TFs regulating these two subclusters with scRNA-seq subcluster 3, SFRP2+WIF1+ fibroblasts, composed of both healthy and SSc cells (Figure S13). Clustering of the TFs from this analysis showed several of the same TFs predicted as driving subcluster 4 differentiation (FOSL2, FOXP1, RUNX2, RUNX1, IRF7), as well as several other TFs (Figure S14). We observed similar TFs if first filtering the inputted genes requiring 6 UMI and expression in 1% of cells (Figure S15). In addition, to correct for potential overfitting due to the variable number of cells per cluster, we downsampled subclusters 3 and 4 40 times and iteratively analyzed predicted TFs by SCENIC. Regulons, regulated in 38 of 40 downsamplings and IRF7, STAT1 and CREB3L1, further validated these TFs in myofibroblast differentiation (Figure 8F).

Surprisingly, none of the transcriptomes in the initial analysis including all transcriptome genes associated with each subcluster identified SMAD2 or SMAD3 regulons, the canonical TFs associated with TGF-β activation. However, if we selected 984 differentially regulated genes between SSc SFRP4+ myofibroblasts, the remaining SSc SFRP2^hi^ cells and the control SFRP2^hi^ fibroblasts, SCENIC detected the SMAD3 regulon. This analysis showed upregulated regulons in a graded fashion, higher in SFRP4^hi^PRSS23+ subcluster 4 fibroblasts and highest in SFRP4+ myofibroblasts, again with some overlapping TFs seen in the previous analyses, including STAT1, FOSL2, RUNX1 and FOXP1 (Figure S16). In this analysis, the SMAD3 regulon was shown upregulated, but its regulation was only associated with the transition between SFRP2^hi^WIF1+ (subcluster 3) to SFRP2^hi^WIF-(subcluster 4) fibroblasts and was actually decreased in SFRP2^hi^SFRP4+, myofibroblasts (Figure S16).

Since SCENIC networks are constructed from the same scRNA-seq dataset they are applied to, we also analyzed predicted TFs based on DoRothEA^55, 56^, which relies on independent TF-targeted gene interactions (regulon activity) curated from various resources, such as the literature, ChIP-seq peaks, TF binding motifs and gene expression inferred interactions. Based on the level of supporting evidence, DoRothEA computed regulons showed several TFs seen on the SCENIC analyses, most consistently SMAD3, STAT1, FOSL2 and HIF1A regulons, upregulated in SFRP2^hi^WIF-(PRSS23+) fibroblasts in most interaction confidence levels, including level A, the level associated with the highest confidence interactions (Figure S17).

As previous studies have strongly implicated TGF-β in SSc pathogenesis^5, 51^, we further validated the role of SMAD3, its downstream TF, on SCENIC predicted target genes in the transcriptome of SSc fibroblasts. To this end, we compared the genes included in the SCENIC, SFRP2^hi^WIF-(PRSS23+), SMAD3 regulon with genes consistently upregulated by TGF-β1-, TGF-β2- or TGF-β3-treated dermal fibroblasts (Table S9). TGF-β induced expression of CHAC1, a SCENIC-predicted downstream target of SMAD3 was inhibited by SIS3, a specific inhibitor of Smad3 phosphorylation^57^ in dermal fibroblasts from both control and SSc subjects (Figure S18).

Because SMAD3 was expressed at low levels (in only a fraction of the cells), regulation of other SMAD3 targets predicted by SCENIC were difficult to assay, even by RT-PCR. Thus, to gain further insight into the role of SMAD3, we developed a SMAD3 activity index based on DoRothEA target A genes. Cells in cluster 4 (SSc SFRP2+ fibroblasts) and some cells also in cluster 3, 0 and 8 (dermal sheath) were found to express higher SMAD3 activity scores (Figure S19).

To further examine SMAD3 activity, we created SMAD3 activity indices from experimentally determined RNA expression after SMAD3 knockdown in myofibroblasts. SMAD3 siRNA depressed SMAD3 expression to 6.7% of non-targeting siRNA treatment (see Table S10). These activity indices showed increased SMAD3 regulon activity in a more restrictive pattern, the highest activity was in cluster 4 (SSc SFRP2+ fibroblasts) and enhanced in the region of the SFPR2+, SFRP4+ myofibroblasts, as well as in cluster 8 (dermal sheath cells, Figure S20).

## DISCUSSION

We show here through bioinformatics and co-staining methods that myofibroblasts in SSc skin are a subpopulation of SFRP2^hi^-expressing fibroblasts. We have shown previously that these cells represent the most common fibroblast population in the skin, with a long narrow morphology that is similar to the morphology seen on staining SSc myofibroblasts with SMA^3^. Bioinformatics analyses show that SFRP2^hi^ and myofibroblasts share closely related transcriptomes and pseudotime analysis indicates that SSc myofibroblasts derive from SFRP2^hi^PRSS23+WIF1− fibroblasts, an SFRP2^hi^ fibroblast subpopulation. Our human SSc data is consistent with murine data, showing that DPP4-expressing fibroblasts in mice are profibrotic in wound healing^26^. However, DPP4, along with SFRP2, mark the largest population of fibroblasts in human dermis^27^, and our scRNA-seq data provide more specific markers for this and related fibroblast subpopulations. The Rinkevich et al cell lineage tracing study strongly supports the pseudotime analysis of our data, showing that the SFRP2/DPP4 fibroblast subpopulation in healthy skin is the progenitor of fibrogenic fibroblasts in SSc skin, including both SFRP2^hi^PRSS23+WIF1− fibroblasts and myofibroblasts^26^.

Our data show a global shift in fibroblast phenotypes in SSc skin. This includes increased expression of PRSS23 and other genes by SFRP2^hi^ fibroblasts, but also a shift within the population of APOE^hi^/CCL19/C7 fibroblasts, which show strikingly upregulated expression of a distinct series of genes like CCL19 not upregulated in SSc SFRP2^hi^ fibroblasts. These observations indicate that SSc is not a disease affecting only myofibroblasts. On the other hand, many genes, such as TNC, are regulated across different fibroblast subpopulations in SSc skin, suggesting that these different fibroblasts are being exposed to a common stimulus, such as Wnt or TGF-β.

Fibroblasts in SSc skin differentiate into myofibroblasts in two steps. The first step, a global shift of SFRP^hi^WIF1+ fibroblasts to SFRP2^hi^PRSS23+WIF1− fibroblasts, is likely parallel to that described in Nazari et al, which was mimicked in vitro by inflammatory stimuli (TNFα and LTβ), but not TGF-β^36^. However, some of the key genes upregulated in this first step, such as TNC and THBS1, are known TGF-β responsive genes, and both SCENIC and DoRothEA predicted SMAD3 as regulating the transcriptome of cell in this step. Other data more strongly support TGF-β as driving the second step, transition of SFRP2^hi^PRSS23+WIF1− fibroblasts to myofibroblasts, as many of the genes upregulated in myofibroblasts are known TGF-β targets and are in a cluster of genes downregulated in the skin of SSc patients after treatment with anti-TGF-β/fresolimumab treatment of SSc patients, such as THBS1, COMP, SERPINE1, COL10A1, CTGF, MATN3^51^. Data mapping genes from SMAD3 knockdown experiments supported the role of SMAD3 in both of these steps.

In contrast to work showing DPP4 fibroblasts as profibrotic^26^, other murine studies have shown that adipocytes^35^, pericytes^34^, and myeloid cells^58, 59^ or combinations of these in addition to resident fibroblasts^33^ can act as myofibroblast progenitors. Lineage tracing experiments have elegantly shown that adipocytes at the interface with the dermis contribute to myofibroblasts found upon bleomycin-induced skin fibrosis^35^. Based on observations in patients with less severe myofibroblast infiltration, myofibroblasts appear first at the interface between subcutaneous fat and reticular dermis^3^. Notably, in vitro, TGF-β induces SFRP2, TNC and CTGF expression by adipocytes differentiated in vitro from human adipose-derived progenitors, suggesting that adipocytes might differentiate into SFRP2^hi^PRSS23+WIF1− fibroblast, myofibroblast progenitors. Despite the proximity of SSc myofibroblasts to fat, we did not see any transcriptome relationship or overlap in specific markers between preadipocytes and myofibroblasts. Another study has shown that PDGFRA/PDPN, ADAM12-expressing perivascular cells are progenitors of myofibroblasts in murine skin and neural injury^34^. We found ADAM12 expression highly induced in SSc myofibroblasts, these cells also expressing PDGFRA and PDPN. This contrasts to ADAM12-expressing perivascular progenitors, which downregulate ADAM12 during myofibroblast differentiation. In addition, we did not see co-expression of pericyte markers to suggest myofibroblast differentiation from a pericyte progenitor, and pericyte populations did not significantly express ADAM12 (see Table S3). Although we cannot exclude the possibility that SSc myofibroblasts differentiate from a non-pericyte perivascular progenitor, our pseudotime analysis as well as SSc myofibroblast morphology and topological location indicates that they differentiate from SFRP2^hi^ fibroblasts, cells that are distributed throughout normal dermis. Another recent scRNA-seq study identified myeloid cell markers *CD45* and *LYZ2* in a subcluster of wound myofibroblasts^60^. However, we did not observe expression of any myeloid marker genes in SSc SFRP2^hi^PRSS23+WIF1− or SSc myofibroblasts (Table S7, and data not shown). In contrast to these cell types in which we could find little transcriptome evidence for a progenitor relationship, we found multiple genes shared between dermal sheath cells and myofibroblasts, including high ACTA2 expression, the gene encoding SMA^37^. Dermal sheath cells express other markers common to myofibroblasts including COL11A1, and cluster proximal to myofibroblasts in UMAP plots. Despite these similarities, dermal sheath cells do not appear to be the direct progenitors of myofibroblasts in SSc skin, as each cell type expresses distinct sets of genes, and SSc fibroblasts are transcriptionally and topologically much more closely related to SFRP2^hi^PRSS23+WIF1− fibroblasts.

We also show increased proliferation of SFRP2^hi^PRSS23+WIF1− fibroblasts in SSc skin. Although, this low rate of proliferation is unlikely to account for the appearance of this cell type in SSc skin, it does suggest that a fibroblast growth factor contributes to the altered phenotype of these cells.

Several genes regulated in SSc myofibroblasts are reciprocally regulated compared to SM22-promoter tdTomato sorted wound myofibroblasts as the wound heals and the fibroblasts lose SMA expression (^61^, Table S6): TNC, SERPINE2, IGFBP3 (increased in SSc and early SMA+ wound myofibroblasts), WIF1 (decreased in SSc and late SMA+ wound myofibroblasts). Despite these parallels most gene expression changes seen in SSc myofibroblasts are distinct from those seen in wound myofibroblasts. This may have to do with the limitation of selecting myofibroblasts based on the SM22 promoter, as TAGLN (the target of the SM22 promoter) is also expressed by smooth muscle cells^62^ and we see TAGLN (the target of the SM22 promoter) also highly expressed by pericytes and dermal sheath cells (not shown). Alternatively, wound and SSc myofibroblasts may represent different cell types and originate from different progenitors.

We originally described altered Wnt pathway gene expression in skin fibrosis, showing that Wnt2 and SFRP4 mRNAs are strongly upregulated in the Tsk murine model of skin fibrosis, as well as in SSc skin biopsies^38^. Subsequently in a more comprehensive analysis of Wnt-related genes, we confirmed upregulated and correlated expression of WNT2 and SFRP4 gene expression in SSc skin^63^. We show here that the correlated upregulation of WNT2 and SFRP4 expression in SSc skin is most likely due to their co-regulation in SSc myofibroblasts. We also showed previously that SSc skin shows markedly decreased expression of WIF1 (−7.88-fold), a soluble Wnt inhibitor^63^. Subsequent work by others confirmed down-regulation of WIF1 and increased Wnt activity^64^. Our data here show that downregulated WIF1 is a marker for a global shift in the phenotype of SFRP2hi-expressing fibroblasts as they transition from SFRP2hiWIF1+ to SFRP2hiPRSS23+WIF1− fibroblasts. As we have previously shown that WIF1 expression in whole skin biopsies correlates strongly inversely with the MRSS^42^, this supports the importance of this global shift in fibroblast phenotype that appears to precede the differentiation of these cells into myofibroblasts.

Several studies have implicated Wnts in fibrosis^65^. Wnt pathway activation increases both fibrillin matrix^63^ and collagen expression. Wnt10b or β-catenin overexpression in mice leads to dermal fibrosis with increased expression of COL1A1, COL1A2, CTGF and ACTA2 mRNA in the skin^66, 67^. Wnt3a blocks preadipocyte differentiation into adipocytes and stimulates their differentiation into myofibroblasts ^64^. Other studies indicate that TGF-β mediates fibrosis by inhibiting DKK1, an endogenous Wnt inhibitor, leading to unrestrained profibrotic Wnt activity^68^. Although Wnt10b and DKK1 up- and down-regulation, respectively, have been identified by IHC^66, 68^, our microarray of whole skin shows Wnt10b expression decreased and DKK1 DKK2 and DKK3 all increased in SSc compared to control skin (combined microarray data; R. Lafyatis). Thus, if indeed these Wnts are playing key roles in SSc pathogenesis, then there is a disconnect between mRNA and protein expression of these genes. This is not an unusual occurrence and indeed a significant limitation to gene expression analyses. However, we propose that altered expression of WIF1, SFRP4 and WNT2, all of whose expression correlates highly with the MRSS, are more likely the key deregulated Wnts in SSc skin.

Regulon analysis implicated several unexpected TFs in regulating the transcriptome of SSc fibroblast differentiation, particularly STAT1, FOSL2, RUNX1, IRF7, HIF1A, CREB3L1 and FOXP1, as well as SMAD3. IRF7 is upstream^69^ and STAT1 downstream^70, 71^interferon signaling, previously implicated in SSc skin^41^. Polymorphisms in the IRF7 and HIF1A genes are associated with SSc^72, 73^; IRF7 can bind SMAD3, and regulate fibrosis and profibrotic gene expression^74^, while HIF1A has been implicated in mediating hypoxia induced skin fibrosis^75^. Transgenic FOSL2 overexpression leads to murine skin fibrosis, reproducing several features of SSc^76^. FOSL2 is induced by TGF-β and regulates collagen production. Thus, several of the TFs predicted to regulate SSc fibroblast differentiation have been implicated in SSc skin fibrosis.

In conclusion, we identify the transcriptome of SSc myofibroblasts and show that SFRP4 is an immunohistochemical marker for these cells. Further, our bioinformatics analyses indicate that myofibroblasts differentiate in a two-step process from SFRP4/DPP4-expressing normal fibroblast progenitors. These data also provide direct insights into previous studies of altered gene expression in SSc skin. We anticipate that applying scRNA-seq in clinical trial settings will enable far greater insights into the effects of therapeutics on the complex alterations of various cell types in SSc skin. We also expect that these observations will provide insights into myofibroblast origin and differentiation in other fibrotic diseases.

## METHODS

### Study Approval

The University of Pittsburgh Medical Center Institutional Review Board (Pittsburgh, PA, USA) reviewed and approved the conduct of this study. Written informed consent was received from all participants prior to inclusion in the study.

### Single cell RNA-sequencing

3 mm skin biopsies were obtained from study subjects, digested enzymatically (Miltenyi Biotec Whole Skin Dissociation Kit, human) for 2 hours and further dispersed using the Miltenyi gentleMACS Octo Dissociator. The resulting cell suspensions were filtered through 70 micron cell strainers twice and re-suspended in PBS containing 0.04% BSA. Resulting cell suspensions were loaded into 10X Genomics Chromium instrument (Pleasanton, CA) for library preparation as described previously ^27^. V1 and V2 single cell chemistries were used per manufacturer’s protocol. Libraries were sequenced (~200 million reads/sample), using the Illumina NextSeq-500 platform. The sequencing reads were examined by quality metrics, transcripts mapped to reference human genome (GRCh38) and assigned to individual cells according to cell barcodes, using Cell Ranger (10X Genomics).

Data analysis was performed using R (version 3.5). Seurat 3.0 was used for data analysis, normalization of gene expression, and identification and visualization of cell populations ^77, 78^, largely as described previously ^27^. Cell populations were identified based on gene markers and visualized by t-distributed stochastic neighbor embedding (t-SNE) ^79^ or UMAP ^80^ plots. We used AddModuleScore to calculate the average expression levels of each program (cluster) on a single cell level, subtracted by the aggregated expression of control feature sets. Pathway analysis was performed with Gene Ontology Enrichment Analysis. Data presented were normalized between samples using SCTransform which models technical noise using a regularized negative binomial regression model ^81^.

### Immunohistochemistry and Imaging

Tyramide SuperBoost kit (Invitrogen, USA) was used to amplify signals in co-stained tissues as per manufacturers protocol. Briefly, IF of formalin-fixed, paraffin-embedded human forearm skin biopsies were first, deparaffinized and rehydrated for antibody staining. Slides were placed in citrate buffer pH 6, steamed for 20 minutes and cooled 20 minutes at room temperature for heat induced antigen retrieval before washing in phosphate buffered saline. All primary antibodies were incubated overnight at 4° Celsius. All poly-HRP secondary antibodies were used as per manufacturers protocol along with tyramide stock solution. Tyramides were incubated for 5 minutes each before neutralized with stop solution. Monoclonal mouse anti-SMA (1:1000; M0851; Clone14A; Dako, Denmark AS, Denmark) was labeled with Alexa Fluor 647 tyramide solution. In order to multiplex staining of slides with antibodies from the same species, slides were placed in citrate buffer pH 6, steamed for 20 minutes and cooled 20 minutes at room temperature for unbound antibody stripping before washing in phosphate buffered saline and proceeding with next antibody. Next, monoclonal mouse SFRP2 (1:250; MAB539; Millipore, USA) labeled with Alexa Fluor 488 tyramide solution was applied and washed, and then polyclonal rabbit SFRP4 (1:500;153287-1-AP; Proteintech, USA) was labeled with Alexa Fluor 568.

For single staining polyclonal rabbit anti-CCL19 (1:500, ab221704, Abcam, USA) was labeled with Alexa Fluor 568 tyramide solution; monoclonal mouse anti-CRABP1 (1:500, MA3-813, C-1, ThermoFisher, USA) was labeled with Alexa Fluor 488 tyramide solution; polyclonal rabbit anti-POSTN (1:250, ab14041, Abcam, USA) was labeled with Alexa Fluor 568 tyramide solution; and monoclonal mouse anti-SLPI (1:50, [31] ab17157, Abcam, USA) was labeled with Alexa Fluor 568 tyramide solution.

All slides were counterstained with nuclear stain Hoechst 33342 (Invitrogen, USA) and cover slipped with Pro-Long™ Diamond Antifade Mountant (P36961: Life Technologies, USA). Images were acquired using an Olympus Fluoview 1000 Confocal Scanning microscope.

### Transcription factor inference-

#### SCENIC

In order to better understand the TFs activating gene expression in SSc fibroblasts and myofibroblasts, we utilized SCENIC ^54^, a computational method for detecting gene regulatory networks. Embedded in this method is the identification of regulons, groups of genes identified by their co-expression with transcription factors (TFs, GENIE3 ^82^), further selected by showing that genes in the regulon are enriched for TF cis-regulatory motifs. SCENIC then scores each cell for the level of gene expression by genes in each regulon, reported as AUC.

For the SCENIC analyses, we used only cells from V2 chemistries, 4 control and 9 SSc samples. To begin clusters 1, 2, 3, and 4 were subsetted from the fibroblast dataset and all genes showing expression in at least one cell were analyzed. A second analysis was then carried out subsetting clusters 3 and 4, with cluster 4 further subsetted to delineate the SFRP4+ myofibroblast group. Again, all genes showing expression in at least one cell were included in the analysis. Alternatively, we filtered cells based on the workflow provided by Aerts lab^54^, keeping genes that 1) with at least 6 UMI across all cells, and 2) detected in at least 1% of cells.

Finally, to focus on changes associated with SSc, a more restrictive gene list (984 genes) was compiled of: 1) genes increased in SFRP2+ SSc cells (in clusters 3 and 4) compared to control SFRP2+ cells (in clusters 3 and 4, Bonferroni corrected Wilcoxon p<0.05); and 2) genes increased in SFRP2+SFRP4+ myofibroblasts compared to SSc SFRP2+SFRP4-cells (in cluster 3 and 4, Bonferroni corrected Wilcoxon p<0.05).

Using SCENIC we then analyzed the single-cell RNA-seq expression matrices by GENIE3 to infer the co-expression network. GENIE3’s output, a link list, included the potential regulators for each gene along with their weights, these weights representing the relevance the TF has in the prediction of the gene target.

#### DoRothEA and VIPER

We used VIPER in combination with DoRothEA to estimate transcription factor (TF) activities from gene expression data^55^. DoRothEA contains 470,711 interactions, covering 1396 TFs targeting 20,238 genes, which rely on the independent TF-targeted gene interactions (regulon activity) curated from various resources such as literature, ChIP-seq peaks, TF binding motifs and gene expression inferred interactions. Based on the number of supporting evidences that accompany each interaction, an interaction confidence level is assigned, ranging from A to E, with A being the highest confidence interactions and E the lowest. VIPER is a statistical method that used in combination with DoRothEA to estimate TF activities from scRNA-seq expression data^83^.

We used TF target genes from DoRothEA, level A and from genes downregulated by siRNA to SMAD3 to create SMAD3 activity scores. Activity scores were also derived from SMAD3 siRNA treated myofibroblasts, filtered for absolute gene expression of non-targeting control siRNA >50 TPM, showing expression of SMAD3 siRNA treated cells of less than 0.8 of non-targeting control siRNA, and excluding genes showing expression less than 0.7 in HRPT1 siRNA treated cells compared to non-targeting control siRNA (415 genes); or for absolute gene expression in non-targeting control treated siRNA cells of >100 TPM, showing expression of SMAD3 siRNA treated cells of less than 0.7 of non-targeting control siRNA, and excluding genes showing expression less than 0.7 in HRPT1 siRNA treated cells compared to non-targeting control siRNA (74 genes).Using the Seurat AddModuleScore, we plotted these activity scores on scRNA-seq UMAP feature plots.

#### Downsampling

In order to test if the difference in cell numbers among clusters of 3 (WIF1+), 4 (PRSS23+), and 4 (SFRP4+) (673, 743, 73 respectively) would affect the SCENIC regulatory analysis, we used R function ‘sample’ to randomly select 73 cells from cluster 3_WIF1+ and 4_PRSS23+ and then performing SCENIC analysis. We downsampled and performed SCENIC analyses 40 times with the resulting, different pools of cells.

### Pseudotime analysis

Expression values were normalized in Monocle 3 (accounting for technical variation in RNA recovery as well as sequencing depth) by estimating size factors for each cell and the dispersion function for genes ^45, 84^. Nonlinear dimensionality reduction was performed using UMAP. Cells were organized into trajectories by Monocle using reversed graph embedding (a machine learning strategy) to learn tree-like trajectories. Once a principal graph had been learned, each cell was projected onto it, using SimplePPT, the default method in Monocle 3; it assumes that each trajectory is a tree that may have multiple roots.

### RNA velocity analysis

Velocyto, a package for the analysis of expression dynamics in single-cell RNA seq data based on spliced and unspliced transcript reads, was used to estimate the time derivative of the gene expression state ^46^. RNA velocities of cells in clusters 1,3 and 4 were estimated using gene-relative model with k-nearest neighbor cell pooling (k=50) based on top 100 differentially expressed genes from myofibroblast, cluster 3, and cluster 4. Velocity fields were projected into a UMAP-based embedding through SeuratWrappers in Seurat.

### Microarray Analyses

We combined and clustered microarray data from our previous biomarker study and clinical trials conducted by our center using the same Affymetrix U133A2.0 microarray chips ^42, 51, 52, 53^. Data were normalized using the MAS 5.0 algorithm, gene expression values were clustered using Cluster 3.0 ^85^. After filtering for genes showing differences of greater than 100 across all samples, genes were mean centered, normalized, hierarchically clustered by complete linkage, and visualized using Java Treeview ^86^. Hierarchical clusters (groups of genes referred to as signatures), corresponding to genes associated with the transition of healthy SFRP2+ fibroblasts into SSc SFRP2+ fibroblasts (PRSS23, TNC, THBS1) and into myofibroblasts (CTGF, ADAM12, COL10A1 and MATN3) were analyzed, using the Seurat AddModuleScore function. The Seurat AddModuleScore function calculates the difference between the average expression levels of each gene set compared to random control genes at a single cell level. These values were then plotted on t-SNE feature plots.

### Cell culture, and TGF-β, phospho-SMAD3 and siRNA analyses

Early passage human dermal fibroblasts from SSc or healthy control skin were cultured in DMEM supplemented with 10% FBS after collagenase digestion. Fibroblasts were passaged at ~80% confluence, the following day placed in 0.1% serum and treated with TGF-β1, TGF-β2 or TGF-β3 (R&D Systems) or left untreated (control), or pretreated one hour with the Smad3 phosphorylation inhibitor SIS3 (CAS 1009104-85-1, Sigma Aldrich). After 16 hours, RNA was prepared and analyzed by microarray Affymetrix U133A2.0 microarray chips as above, or cDNA prepared and analyzed by RT-PCR using primers to CHAC1 Taqman FAM-MGB dye (Thermo fisher #4331182) or SMAD3 Taqman FAM-MGB dye (Thermo fisher #4453320). Ct values for each treatment were normalized to 18S. Delta Ct for sample was then normalized to the control treatment and Fold change-calculated.

For siRNA experiments, pulmonary myofibroblasts (passage 6) isolated from lung explants were used in knockdown experiments. Cells were passaged at 60% confluence, washed and then siRNAs targeting SMAD3, HRPT1 (control) or non-targeting control transfected 8 hours using Lullaby Transfection Buffer (OZ Biosciences; LL71000) in 200μL OptiMEM with 5μL 10μM reconstituted dsiRNAs (TriFECTa DsiRNA kit, cat#: hs.Ri.SMAD3.13, Integrated DNA Technology). After 48 hours RNA was isolated from the cells with the RNeasy kit (Qiagen), quantified and quality checked in TapeStation High Sensitivity RNA Screen Tape. cDNA libraries were synthesized and 25 million single-end reads sequenced per sample using an Illumina High-Throughput Sequencer. FastQC reports were used to ensure quality data was entered into analysis. Alignment and Gene Counts were carried out using CLC Genomics version 20.0.3. Transcripts Per Kilobase Million (TPM) were exported for samples and log fold changes were calculated with respect to the non-targeting controls.

### Statistics

For Tables examining differential gene expression between cells or groups of cells within clusters, cells were filtered out that were expressed in less than 10% of the cells showing upregulated expression. Comparisons of average numbers of cells in each fibroblast subpopulation were compared using Wilcoxon Rank-Sum test. Differential gene expression between healthy controls and SSc was assessed using Seurat’s implementation of the non-parametric Wilcoxon rank sum test. A Bonferroni correction was applied to the results. Differences between the average proportions of cells in all control and SSc clusters were compared using the chi square test. Individual differences between proportions of cells in each patient comparing SSc with controls cluster were calculated using Mann-Whitney. All statistical tests were two-sided.

## Data Availability

All scRNA-seq data including Gene cell UMI matrix and a BAM file containing aligned reads are available at the Gene Expression Omnibus: <u>GSE138669</u>.

## Code Availability

Code for data analyses are available and referenced in the text.

## AUTHOR CONTRIBUTIONS

T.T, N.M. M.B, R.B. and A.P. contributed to designing research studies, conducting experiments, acquiring data, analyzing data, and writing the manuscript.

W.C. contributed to analyzing data, and writing the manuscript.

R.D. contributed to designing research studies, conducting experiments, writing the manuscript.

R.L. contributed to designing research studies, analyzing data, and writing the manuscript.

## BIBLIOGRAPHY

1. Man A, Correa JK, Ziemek J, Simms RW, Felson DT, Lafyatis R. Development and validation of a patient-reported outcome instrument for skin involvement in patients with systemic sclerosis. Annals of the rheumatic diseases 76, 1374–1380 (2017).

2. Ziemek J, Man A, Hinchcliff M, Varga J, Simms RW, Lafyatis R. The relationship between skin symptoms and the scleroderma modification of the health assessment questionnaire, the modified Rodnan skin score, and skin pathology in patients with systemic sclerosis. Rheumatology 55, 911–917 (2016).

3. Kissin EY, Merkel PA, Lafyatis R. Myofibroblasts and hyalinized collagen as markers of skin disease in systemic sclerosis. Arthritis Rheum 54, 3655–3660 (2006).

4. Hinz B. Myofibroblasts. Experimental eye research 142, 56–70 (2016).

5. Lafyatis R. Transforming growth factor beta--at the centre of systemic sclerosis. Nature reviews 10, 706–719 (2014).

6. Hinz B. The extracellular matrix and transforming growth factor-beta1: Tale of a strained relationship. Matrix biology: journal of the International Society for Matrix Biology 47, 54–65 (2015).

7. Hinz B. Formation and function of the myofibroblast during tissue repair. J Invest Dermatol 127, 526–537 (2007).

8. Duscher D, et al. Mechanotransduction and fibrosis. Journal of biomechanics 47, 1997–2005 (2014).

9. Huang X, et al. Matrix stiffness-induced myofibroblast differentiation is mediated by intrinsic mechanotransduction. American journal of respiratory cell and molecular biology 47, 340–348 (2012).

10. Pakshir P, Hinz B. The big five in fibrosis: Macrophages, myofibroblasts, matrix, mechanics, and miscommunication. Matrix biology: journal of the International Society for Matrix Biology, (2018).

11. Schulz JN, Plomann M, Sengle G, Gullberg D, Krieg T, Eckes B. New developments on skin fibrosis - Essential signals emanating from the extracellular matrix for the control of myofibroblasts. Matrix biology: journal of the International Society for Matrix Biology, (2018).

12. Wipff PJ, Rifkin DB, Meister JJ, Hinz B. Myofibroblast contraction activates latent TGF-beta1 from the extracellular matrix. The Journal of cell biology 179, 1311–1323 (2007).

13. Klingberg F, et al. Prestress in the extracellular matrix sensitizes latent TGF-beta1 for activation. The Journal of cell biology 207, 283–297 (2014).

14. Tsai CC, Wu SB, Kau HC, Wei YH. Essential role of connective tissue growth factor (CTGF) in transforming growth factor-beta1 (TGF-beta1)-induced myofibroblast transdifferentiation from Graves’ orbital fibroblasts. Scientific reports 8, 7276 (2018).

15. Folger PA, Zekaria D, Grotendorst G, Masur SK. Transforming growth factor-beta-stimulated connective tissue growth factor expression during corneal myofibroblast differentiation. Investigative ophthalmology & visual science 42, 2534–2541 (2001).

16. Shi-Wen X, et al. Endothelin-1 promotes myofibroblast induction through the ETA receptor via a rac/phosphoinositide 3-kinase/Akt-dependent pathway and is essential for the enhanced contractile phenotype of fibrotic fibroblasts. Molecular biology of the cell 15, 2707–2719 (2004).

17. Tang WW, et al. Platelet-derived growth factor-BB induces renal tubulointerstitial myofibroblast formation and tubulointerstitial fibrosis. The American journal of pathology 148, 1169–1180 (1996).

18. Kaur H, et al. Corneal stroma PDGF blockade and myofibroblast development. Experimental eye research 88, 960–965 (2009).

19. Bostrom H, et al. PDGF-A signaling is a critical event in lung alveolar myofibroblast development and alveogenesis. Cell 85, 863–873 (1996).

20. Akasaka Y, et al. Basic fibroblast growth factor in an artificial dermis promotes apoptosis and inhibits expression of alpha-smooth muscle actin, leading to reduction of wound contraction. Wound repair and regeneration: official publication of the Wound Healing Society [and] the European Tissue Repair Society 15, 378–389 (2007).

21. Cushing MC, Mariner PD, Liao JT, Sims EA, Anseth KS. Fibroblast growth factor represses Smad-mediated myofibroblast activation in aortic valvular interstitial cells. FASEB journal: official publication of the Federation of American Societies for Experimental Biology 22, 1769–1777 (2008).

22. Oldroyd SD, Thomas GL, Gabbiani G, El Nahas AM. Interferon-gamma inhibits experimental renal fibrosis. Kidney international 56, 2116–2127 (1999).

23. Nevers T, et al. Th1 effector T cells selectively orchestrate cardiac fibrosis in nonischemic heart failure. The Journal of experimental medicine 214, 3311–3329 (2017).

24. Driskell RR, et al. Distinct fibroblast lineages determine dermal architecture in skin development and repair. Nature 504, 277–281 (2013).

25. Driskell RR, Watt FM. Understanding fibroblast heterogeneity in the skin. Trends in cell biology 25, 92–99 (2015).

26. Rinkevich Y, et al. Skin fibrosis. Identification and isolation of a dermal lineage with intrinsic fibrogenic potential. Science 348, aaa2151 (2015).

27. Tabib T, Morse C, Wang T, Chen W, Lafyatis R. SFRP2/DPP4 and FMO1/LSP1 Define Major Fibroblast Populations in Human Skin. J Invest Dermatol 138, 802–810 (2018).

28. Philippeos C, et al. Spatial and Single-Cell Transcriptional Profiling Identifies Functionally Distinct Human Dermal Fibroblast Subpopulations. J Invest Dermatol 138, 811–825 (2018).

29. Hinz B, Phan SH, Thannickal VJ, Galli A, Bochaton-Piallat ML, Gabbiani G. The myofibroblast: one function, multiple origins. The American journal of pathology 170, 1807–1816 (2007).

30. Paquet-Fifield S, et al. A role for pericytes as microenvironmental regulators of human skin tissue regeneration. The Journal of clinical investigation 119, 2795–2806 (2009).

31. Hinz B, Pittet P, Smith-Clerc J, Chaponnier C, Meister JJ. Myofibroblast development is characterized by specific cell-cell adherens junctions. Molecular biology of the cell 15, 4310–4320 (2004).

32. Wu M, et al. Identification of cadherin 11 as a mediator of dermal fibrosis and possible role in systemic sclerosis. Arthritis & rheumatology 66, 1010–1021 (2014).

33. LeBleu VS, et al. Origin and function of myofibroblasts in kidney fibrosis. Nature medicine 19, 1047–1053 (2013).

34. Dulauroy S, Di Carlo SE, Langa F, Eberl G, Peduto L. Lineage tracing and genetic ablation of ADAM12(+) perivascular cells identify a major source of profibrotic cells during acute tissue injury. Nature medicine 18, 1262–1270 (2012).

35. Marangoni RG, et al. Myofibroblasts in murine cutaneous fibrosis originate from adiponectin-positive intradermal progenitors. Arthritis & rheumatology 67, 1062–1073 (2015).

36. Nazari B, et al. Altered Dermal Fibroblasts in Systemic Sclerosis Display Podoplanin and CD90. The American journal of pathology, (2016).

37. Jahoda CA, Reynolds AJ, Chaponnier C, Forester JC, Gabbiani G. Smooth muscle alpha-actin is a marker for hair follicle dermis in vivo and in vitro. Journal of cell science 99 (Pt 3), 627–636 (1991).

38. Bayle J, Fitch J, Jacobsen K, Kumar R, Lafyatis R, Lemaire R. Increased expression of Wnt2 and SFRP4 in Tsk mouse skin: role of Wnt signaling in altered dermal fibrillin deposition and systemic sclerosis. J Invest Dermatol 128, 871–881 (2008).

39. Agabalyan NA, Rosin NL, Rahmani W, Biernaskie J. Hair follicle dermal stem cells and skin-derived precursor cells: Exciting tools for endogenous and exogenous therapies. Experimental dermatology 26, 505–509 (2017).

40. Uhlen M, et al. Proteomics. Tissue-based map of the human proteome. Science 347, 1260419 (2015).

41. Farina G, Lafyatis D, Lemaire R, Lafyatis R. A four-gene biomarker predicts skin disease in patients with diffuse cutaneous systemic sclerosis. Arthritis Rheum 62, 580–588 (2010).

42. Rice LM, et al. A Longitudinal Biomarker for the Extent of Skin Disease in Patients With Diffuse Cutaneous Systemic Sclerosis. Arthritis & rheumatology 67, 3004–3015 (2015).

43. Bhattacharyya S, et al. Tenascin-C drives persistence of organ fibrosis. Nat Commun 7, 11703 (2016).

44. Greenblatt MB, et al. Interspecies comparison of human and murine scleroderma reveals IL-13 and CCL2 as disease subset-specific targets. The American journal of pathology 180, 1080–1094 (2012).

45. Trapnell C, et al. The dynamics and regulators of cell fate decisions are revealed by pseudotemporal ordering of single cells. Nature biotechnology 32, 381–386 (2014).

46. La Manno G, et al. RNA velocity of single cells. Nature 560, 494–498 (2018).

47. Morse C, et al. Proliferating SPP1/MERTK-expressing macrophages in idiopathic pulmonary fibrosis. The European respiratory journal, (2019).

48. Pendergrass SA, Lemaire R, Francis IP, Mahoney JM, Lafyatis R, Whitfield ML. Intrinsic gene expression subsets of diffuse cutaneous systemic sclerosis are stable in serial skin biopsies. J Invest Dermatol 132, 1363–1373 (2012).

49. Assassi S, et al. Dissecting the heterogeneity of skin gene expression patterns in systemic sclerosis. Arthritis & rheumatology 67, 3016–3026 (2015).

50. Milano A, et al. Molecular subsets in the gene expression signatures of scleroderma skin. PloS one 3, e2696 (2008).

51. Rice LM, et al. Fresolimumab treatment decreases biomarkers and improves clinical symptoms in systemic sclerosis patients. The Journal of clinical investigation 125, 2795–2807 (2015).

52. Lafyatis R, et al. Inhibition of beta-Catenin Signaling in the Skin Rescues Cutaneous Adipogenesis in Systemic Sclerosis: A Randomized, Double-Blind, Placebo-Controlled Trial of C-82. J Invest Dermatol 137, 2473–2483 (2017).

53. Mantero JC, et al. Randomised, double-blind, placebo-controlled trial of IL1-trap, rilonacept, in systemic sclerosis. A phase I/II biomarker trial. Clin Exp Rheumatol 36 Suppl 113, 146–149 (2018).

54. Aibar S, et al. SCENIC: single-cell regulatory network inference and clustering. Nature methods 14, 1083–1086 (2017).

55. Garcia-Alonso L, Holland CH, Ibrahim MM, Turei D, Saez-Rodriguez J. Benchmark and integration of resources for the estimation of human transcription factor activities. Genome Res 29, 1363–1375 (2019).

56. Holland CH, et al. Robustness and applicability of transcription factor and pathway analysis tools on single-cell RNA-seq data. Genome Biol 21, 36 (2020).

57. Jinnin M, Ihn H, Tamaki K. Characterization of SIS3, a novel specific inhibitor of Smad3, and its effect on transforming growth factor-beta1-induced extracellular matrix expression. Mol Pharmacol 69, 597–607 (2006).

58. Fathke C, et al. Contribution of bone marrow-derived cells to skin: collagen deposition and wound repair. Stem cells 22, 812–822 (2004).

59. Ogawa M, LaRue AC, Drake CJ. Hematopoietic origin of fibroblasts/myofibroblasts: Its pathophysiologic implications. Blood 108, 2893–2896 (2006).

60. Guerrero-Juarez CF, et al. Single-cell analysis reveals fibroblast heterogeneity and myeloid-derived adipocyte progenitors in murine skin wounds. Nature communications 10, 650 (2019).

61. Plikus MV, et al. Regeneration of fat cells from myofibroblasts during wound healing. Science 355, 748–752 (2017).

62. Solway J, et al. Structure and expression of a smooth muscle cell-specific gene, SM22 alpha. The Journal of biological chemistry 270, 13460–13469 (1995).

63. Lemaire R, et al. Antagonistic effect of the matricellular signaling protein CCN3 on TGF-beta- and Wnt-mediated fibrillinogenesis in systemic sclerosis and Marfan syndrome. J Invest Dermatol 130, 1514–1523 (2010).

64. Wei J, et al. Wnt/beta-catenin signaling is hyperactivated in systemic sclerosis and induces Smad-dependent fibrotic responses in mesenchymal cells. Arthritis Rheum 64, 2734–2745 (2012).

65. Lafyatis R. Connective tissue disease: SSc-fibrosis takes flight with Wingless inhibition. Nature reviews 8, 441–442 (2012).

66. Wei J, et al. Canonical Wnt signaling induces skin fibrosis and subcutaneous lipoatrophy: a novel mouse model for scleroderma? Arthritis Rheum 63, 1707–1717 (2011).

67. Hamburg-Shields E, DiNuoscio GJ, Mullin NK, Lafyatis R, Atit RP. Sustained beta-catenin activity in dermal fibroblasts promotes fibrosis by up-regulating expression of extracellular matrix protein-coding genes. J Pathol 235, 686–697 (2015).

68. Akhmetshina A, et al. Activation of canonical Wnt signalling is required for TGF-beta-mediated fibrosis. Nature communications 3, 735 (2012).

69. Honda K, et al. IRF-7 is the master regulator of type-I interferon-dependent immune responses. Nature 434, 772–777 (2005).

70. Begitt A, et al. STAT1-cooperative DNA binding distinguishes type 1 from type 2 interferon signaling. Nature immunology 15, 168–176 (2014).

71. Dupuis S, et al. Impaired response to interferon-alpha/beta and lethal viral disease in human STAT1 deficiency. Nature genetics 33, 388–391 (2003).

72. Carmona FD, et al. Novel identification of the IRF7 region as an anticentromere autoantibody propensity locus in systemic sclerosis. Annals of the rheumatic diseases 71, 114–119 (2012).

73. Wipff J, et al. Association of hypoxia-inducible factor 1A (HIF1A) gene polymorphisms with systemic sclerosis in a French European Caucasian population. Scand J Rheumatol 38, 291–294 (2009).

74. Wu M, et al. Interferon regulatory factor 7 (IRF7) represents a link between inflammation and fibrosis in the pathogenesis of systemic sclerosis. Annals of the rheumatic diseases 78, 1583–1591 (2019).

75. Distler JH, et al. Hypoxia-induced increase in the production of extracellular matrix proteins in systemic sclerosis. Arthritis Rheum 56, 4203–4215 (2007).

76. Reich N, et al. The transcription factor Fra-2 regulates the production of extracellular matrix in systemic sclerosis. Arthritis Rheum 62, 280–290 (2010).

77. Satija R, Farrell JA, Gennert D, Schier AF, Regev A. Spatial reconstruction of single-cell gene expression data. Nature biotechnology 33, 495–502 (2015).

78. Stuart T, et al. Comprehensive Integration of Single-Cell Data. Cell 177, 1888–1902 e1821 (2019).

79. van der Maaten L, Hinton G. Visualizing Data using t-SNE. Journal of Machine Learning Research 9, 2579–2605 (2008).

80. Becht E, et al. Dimensionality reduction for visualizing single-cell data using UMAP. Nat Biotechnol, (2018).

81. Hafemeister C, Satija R. Normalization and variance stabilization of single-cell RNA-seq data using regularized negative binomial regression. bioRxiv, (2019).

82. Huynh-Thu VA, Irrthum A, Wehenkel L, Geurts P. Inferring regulatory networks from expression data using tree-based methods. PloS one 5, (2010).

83. Alvarez MJ, et al. Functional characterization of somatic mutations in cancer using network-based inference of protein activity. Nature genetics 48, 838–847 (2016).

84. Cao J, et al. The single-cell transcriptional landscape of mammalian organogenesis. Nature 566, 496–502 (2019).

85. Eisen MB, Spellman PT, Brown PO, Botstein D. Cluster analysis and display of genome-wide expression patterns. Proceedings of the National Academy of Sciences of the United States of America 95, 14863–14868 (1998).

86. Saldanha AJ. Java Treeview--extensible visualization of microarray data. Bioinformatics 20, 3246–3248 (2004).

